# Stromal inflammation is a targetable driver of hematopoietic aging

**DOI:** 10.1101/2021.03.08.434485

**Authors:** Evgenia V. Verovskaya, Carl A. Mitchell, Fernando J. Calero-Nieto, Aurélie Hérault, Paul V. Dellorusso, Xiaonan Wang, Si Yi Zhang, Arthur Flohr Svendsen, Eric M. Pietras, Sietske T. Bakker, Theodore T. Ho, Berthold Göttgens, Emmanuelle Passegué

**Affiliations:** Columbia Stem Cell Initiative, Department of Genetics and Development, Columbia University Irving Medical Center, New York, New York 10032, USA; The Eli and Edythe Broad Center of Regeneration Medicine and Stem Cell Research, Department of Medicine, Division Hematology/Oncology, University of California San Francisco, San Francisco, California 94143, USA; Welcome and MRC Cambridge Stem Cell Institute, Department of Hematology, Cambridge University, Jeffrey Cheah Biomedical Centre Puddicombe Way, Cambridge CB2 0AW, UK

**Author notes:** Corresponding author: Emmanuelle Passegué, PhD; Columbia Stem Cell Initiative, Department of Genetics and Development, Columbia University Irving Medical Center, New York, New York 10032, USA. Equal contributions (Evgenia V. Verovskaya, Carl A. Mitchell).

## Abstract

Hematopoietic aging is marked by a loss of regenerative capacity and skewed differentiation from hematopoietic stem cells (HSC) leading to dysfunctional blood production. Signals from the bone marrow (BM) niche dynamically tailor hematopoiesis, but the effect of aging on the niche microenvironment and the contribution of the aging niche to blood aging still remains unclear. Here, we characterize the inflammatory milieu in the aged marrow cavity that drives both stromal and hematopoietic remodeling. We find decreased numbers and functionality of osteogenic mesenchymal stromal cells (MSC) at the endosteum and expansion of pro-inflammatory perisinusoidal MSCs with deterioration of sinusoidal endothelium in the central marrow, which together create a degraded and inflamed old niche. Molecular mapping at single cell resolution confirms disruption of cell identities and enrichment of inflammatory response genes in niche populations. Niche inflammation, in turn, drives chronic activation of emergency myelopoiesis pathways in old HSCs and multipotent progenitors (MPP), which promotes myeloid differentiation at the expense of lymphoid and erythroid commitment and hinders hematopoietic regeneration. Remarkably, niche deterioration, HSC dysfunction and defective hematopoietic regeneration, can be improved by blocking inflammatory IL-1 signaling. Our results demonstrate that targeting niche inflammation is a tractable strategy to restore blood production during aging.

---

Hematopoietic function severely declines with age, causing anemia, impaired adaptive immunity, cancer and autoimmune disease in the elderly^1^. Aging also leads to major changes in bones and the marrow cavity, which together provide specialized niches for HSCs and hematopoietic progenitors^2–4^. Blood production is tailored in part by differential production of distinct HSC-derived MPP subsets with specific myeloid (MPP2, MPP3) and lymphoid (MPP4) lineage biases, which in turn give rise to an array of lineage-restricted progenitors and mature cells^5,6^. The niche and factors present in the marrow milieu, in particular pro-inflammatory cytokines, play important roles in controlling blood output by regulating the number and activation of HSCs, tuning the production and differentiation of lineage-biased and lineage-restricted progenitors, modulating stress hematopoiesis, and exerting context-dependent roles in leukemic transformation^3,4,6,7^. Circulating levels of pro-inflammatory cytokines increase with age and contribute to the stereotypical decline in tissue integrity with age^8,9^. By mid-life (~ 50 years in humans/12 months in mice), the bones themselves are already old with bone thinning, reduced fracture repair potential, and impaired endocrine function^10^. In contrast, the most salient features of blood aging are not fully pronounced until later in life (~ 70 years in humans/20-24 months in mice) and include expansion of dysfunctional HSCs with reduced engraftment potential and impaired blood production capability^2^. While it is now well appreciated that cell-intrinsic features such as replication stress, epigenetic remodeling and metabolic rewiring, are important drivers of old HSC dysfunction^2^, much less is known about the mechanisms driving niche aging and the influence of the aged niche on hematopoietic aging.

Different cell types like MSCs and their osteoprogenitor (OPr) derivatives, endothelial cells (EC), the parasympathetic nervous system, adipocytes and specific mature blood cells like megakaryocytes (Mk) and macrophages serve as important niche components^3,4^, which have now been extensively imaged and profiled with high resolution^11–13^. The marrow cavity is also partitioned between the bone endosteum and the central marrow perivascular space, with each location exhibiting specific cellular compositions and distinct functions in regulating HSC maintenance, progenitor differentiation and blood production^2–4^. Recent studies have described age-related changes in several niche components, in particular changes in the vasculature^14,15^, inervation^16,17^ and specific stromal populations^18,19^, and reported heightened levels of inflammation in the aged marrow cavity^17,20–22^. However, the molecular underpinnings of stromal aging and the key inflammatory mediators that could be targeted to improve hematopoietic aging have yet to be identified.

## Chronically inflamed old BM milieu

To quantify changes in the marrow milieu with age, we used a 200-plex array to screen cytokines present in BM fluids harvested from young and old C57BL/6 wild type (WT) mice (**Fig. 1a; SI Table 1**). 81 analytes were detected at both ages, with 32 upregulated and 11 downregulated with age. The largest group represented inflammatory factors, some of which have been previously reported to increase with age such as RANTES^20^, Eotaxin (CCL11)^23^, and IL-1^21,24^. The second largest group comprised soluble ligands of endothelial cells like ICAM-1 and E-selectin, whose increased levels are hallmarks of vascular activation and inflammation^25^. Other categories included cytokines and chemokines involved in bone^26^ and fat^27^ production and function, which likely reflect major stroma remodeling with age. Complementary analyses with cytokine bead-array measurements confirmed the pro-inflammatory state of the old BM milieu, with significantly elevated levels of IL-1α/β, MIP1α, and TNFα (**Fig. 1b**). These results reveal dramatic changes in secreted factors and demonstrate localized low-grade inflammation in the old marrow.

**Figure 1.**
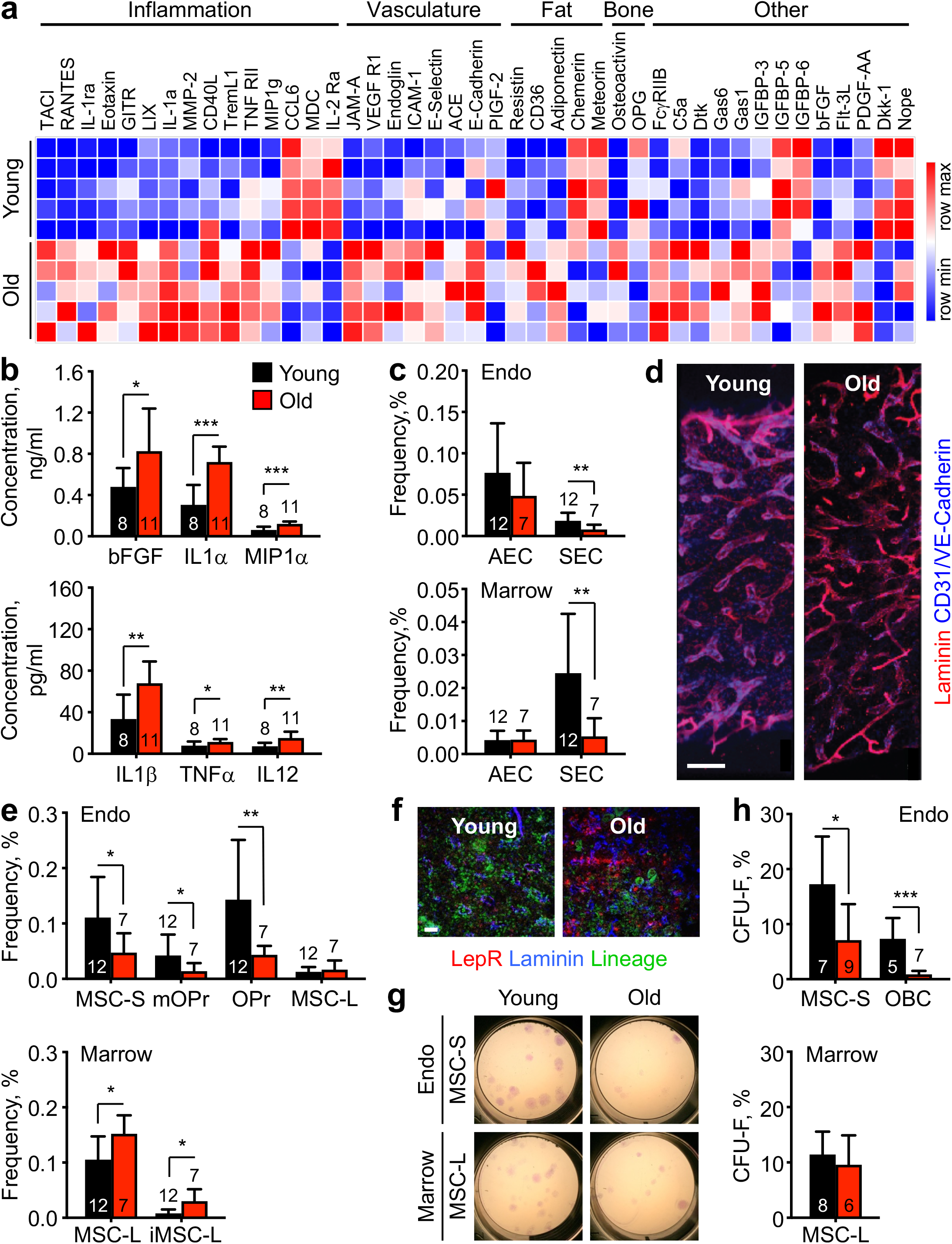
Inflamed milieu and remodeled BM microenvironment with age. **a,** Differentially secreted cytokines in young and old BM fluid measured by 200-plex array and clustered based on broad biological functions (n = 5). **b,** Bead-array measurements of pro-inflammatory cytokines in young and old BM fluids. **c**, Changes in endosteal (Endo) and central marrow (Marrow) EC populations with age. **d,** Whole mount staining of the BM vasculature in young and old mice. Scale bar, 100 μm. **e**, Changes in endosteal and central marrow mesenchymal populations with age. **f,** Representative image of MSC-L immunofluorescence staining in young and old BM. Scale bar, 38 μm. **g-h**, Representative pictures (g) and quantification (h) of fibroblast colony-forming units (CFU-F) from young and old MSC-S, OBC and MSC-L. Data are means ± S.D; *p ≤ 0.05, **p ≤ 0.01, ***p ≤ 0.001.

## Aging remodels the BM niche

To assess how aging affects the niche, we first performed an overall survey of bone structure and composition in young and old mice. Hematoxylin and eosin (H&E) staining showed infrequent accumulation of adipocytes, and only in certain bones, suggesting that aging in mice is not consistently associated with increased marrow adipogenesis (**Extended Data Fig. 1a**). In contrast, old mice showed a fully penetrant accumulation of Mks, which was accompanied by elevated TGFβ and TPO production (**Extended Data Fig. 1a, b**) and is consistent with the Mk-bias of old HSCs^28^. Micro-computed tomography (μCT) confirmed the pronounced decay of trabecular and cortical bones^29^, which correspond to a reduction in bone-lining ALCAM^+^ osteoblasts (**Extended Data Fig. 1c, d**).

We next employed a refined version of our flow cytometry-based method^30^ to investigate both endothelial and mesenchymal niche populations at the endosteum using crushed bone chips and in the central marrow cavity using flushed marrow plugs (**Extended Data Fig. 2a**). While the overall frequency and preferential endosteal location of arteriolar ECs (AEC) was not affected by age, we observed a significant reduction in sinusoidal EC (SEC) frequency at both locations in old mice (**Fig. 1c**). This contrasts with whole mount immunofluorescence analyses showing enhanced branching and dysmorphia of the aged central marrow sinusoidal network (**Fig. 1d**). As previously described^14,15^, we also found unchanged vascular volume and increased vascular leakiness with age (**Extended Data Fig. 2b, c**), which was associated with decreased endocytosis of injected Dragon Green beads (DGB) in old SECs. These results suggest that old SECs are increasingly fragilized and dysfunctional, and are likely lost during enzymatic digestion for cytometric analyses. Accordingly, age-related changes in vascular stiffness and BM hypoxia have been reported to alter vascular integrity and contribute to degraded endothelial integrity^31^. Old mice also showed significantly decreased frequency of endosteal periarteriolar Sca-1^+^ MSCs (MSC-S) and their osteoblastic lineage cell (OBC) derivatives, multipotent PDGFRα^+^ OPr (mOPr) and more committed PDGFRα^−^ OPr (**Fig. 1e; Extended Data Fig. 2a**). Conversely, we found an increase in central marrow perisinusoidal LepR^+^ MSCs (MSC-L) frequency and the striking emergence of an inflammatory Sca-1^low^ subset of MSC-L (iMSC-L) with age (**Fig. 1e, f; Extended Data Fig. 2a**). Functional assessment revealed a consistent loss of fibroblastic colony-forming units (CFU-F) potential in old endosteal MSC-S and OBC populations, but not from old central marrow MSC-L (**Fig. 2g, h**). Loss and functional deterioration of endosteal mesenchymal populations were already observed in 13 month-old middle-age mice (**Extended Data Fig. 2d**), supporting the idea that niche aging precedes hematopoietic aging. The functional decline of old MSC-S also appeared to be cell-intrinsic since both young and old BM cells could similarly stimulate young MSC-S colony formation, but young BM cells could not rescue the defective growth of old MSC-S in co-culture experiments (**Extended Date Fig. 2e**). Collectively, these results reveal a complex remodeling of the old niche, with loss and functional deterioration of endosteal mesenchymal populations, and expansion of inflammatory MSC-L alongside a fragilization of the sinusoidal network in the central marrow. They also indicate that the expanded MSC-L compartment in old mice does not functionally compensate for age-related bone loss, maybe due to the recently described loss of osteogenic osteolectin-expressing periarteriolar MSC-L subset^19^.

**Figure 2.**
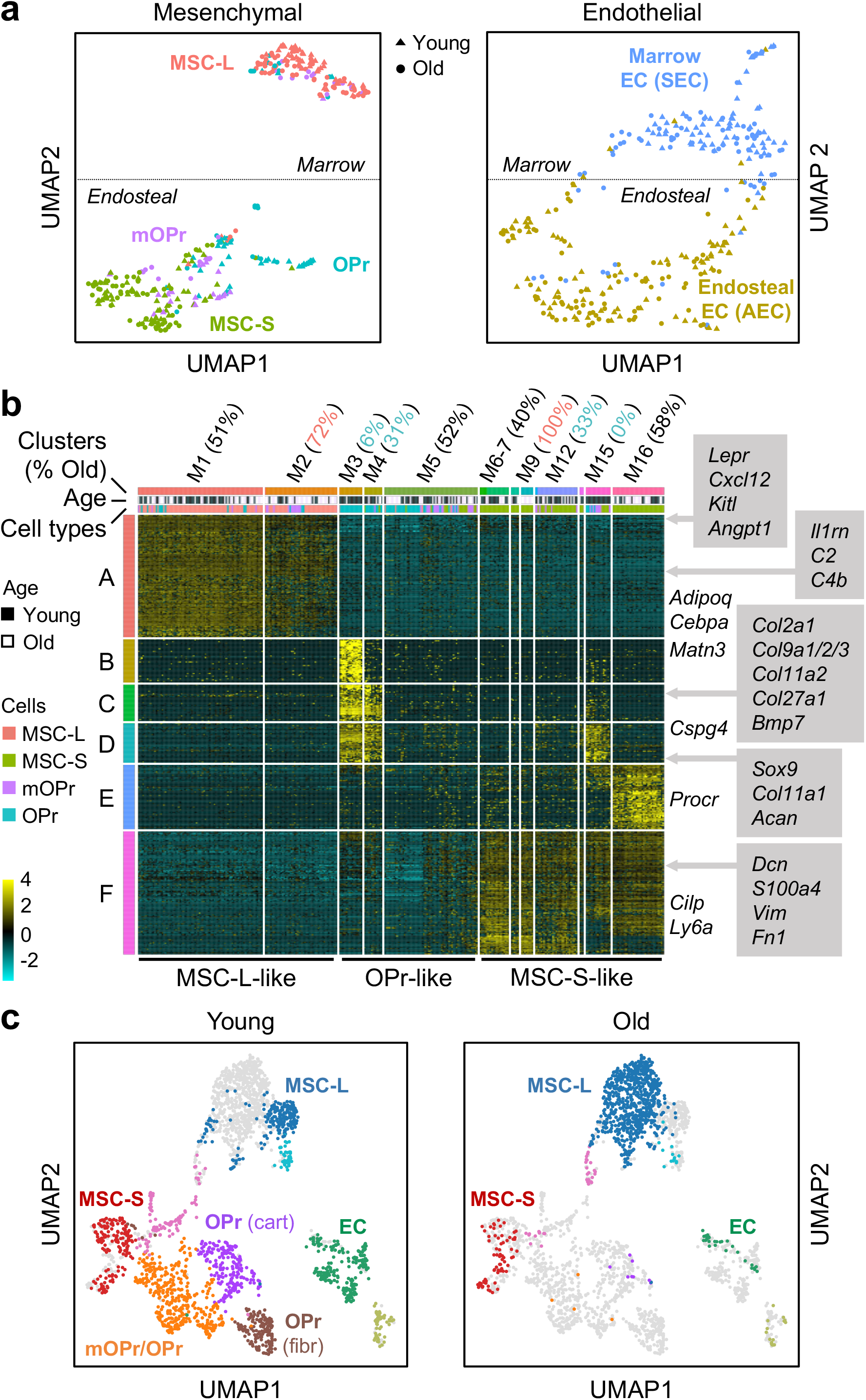
Molecular characterization of the old BM niche. **a,** UMAP visualisation of plate-based scRNAseq analyses of mesenchymal and endothelial populations isolated from young (n=3) and old (n=4) mice. **b**, ICGS output of young and old mesenchymal populations with 16 clusters of cells (M1 to M16) defined according to the expressing pattern of the 6 clusters of genes (A to F). Examples of genes included in gene clusters A to F are shown. **c**, UMAP visualisation of droplet-based scRNAseq analyses of young (n=2) and old (n=1) endosteal and central marrow stromal fractions.

## Altered signaling in old BM niche cells

To understand the molecular changes driving remodeling of the old niche, we performed plate-based single cell RNA sequencing (scRNAseq) analyses on mesenchymal and endothelial populations isolated from both endosteum and central marrow of young and old mice. Uniform manifold approximation and projection (UMAP) representation confirmed preservation of the overall niche structure with age (**Extended Data Fig. 3a**). Further visualization of mesenchymal populations highlighted the distinction between marrow MSC-L and endosteal MSC-S/mOPr/OPr, and also revealed significant contamination of MSC-L by old endosteal populations (**Fig. 2a**). Iterative clustering and guide-gene selection (ICGS) analyses identified 16 unique cell clusters, with M1-2 associated with MSC-L identity, M3-5 with OPr identity, and M6-16 with MSC-S identity (**Fig. 2b**). While most clusters had similar contribution from young and old cells, several contained overwhelmingly old cells (M2 and M9) while others had an under-representation of old cells (M3, M4, M12 and M15). Strikingly, M3/M4/M15 expressed genes specifically associated with osteoblastic and chondrogenic differentiation (**SI Table 2**), supporting a broad loss of bone-forming cells with age. M9 was exclusively composed of old cells with downregulated bone formation pathways, while M2 was dominated by old cells with reduced expression of MSC-L identity genes (**Extended Data Fig. 3b**). In fact, a large fraction of the isolated old endosteal mOPr (45%) and OPr (58%) displayed M1 and M2 cluster identity gene expression, suggesting that they were in fact old endosteal MCS-L likely mis-sorted as OBCs because of reduced LepR surface expression (**Extended Data Fig. 3c**). Categorization of scRNAseq data into MSC-L-like, OPr-like and MSC-S-like gene identity groups showed no major changes in the expression of HSC maintenance factors like *Kitl* and *Cxcl12* between young and old cells, with the numerical expansion of MSC-L cells likely contributing to the elevated SCF levels found in old BM fluids (**Extended Data Fig. 3d**). In contrast, gene set enrichment analysis (GSEA) of differentially expressed genes (DEG) between young and old cells within each identity group highlighted a broad enrichment of inflammatory signaling pathways in all old mesenchymal populations (**Extended Data Fig. 3e; SI Table 3**). Collectively, these molecular analyses demonstrate that the age-related loss of endosteal populations and expansion of marrow MSC-L are even more pronounced than quantified by flow cytometry due to surface marker infidelity in old MSC-L. They also indicate that a loss of osteoblastic/chondrogenic commitment and chronic activation of inflammatory response underpin the functional deterioration of old endosteal populations and associated bone loss. Moreover, they suggest that the expansion of *Kitl*-expressing MSC-L with degraded cell identity serves to support an expanded pool of dysfunctional old HSCs.

Similar UMAP visualization of endothelial populations clearly separated marrow ECs (mostly composed of SECs according to index sorting data) from endosteal ECs (dominated by AECs) regardless of the age of the populations (**Fig. 2a**). ICGS analyses identified 7 cell clusters, with E3-4 associated with AEC identity and E7 with SEC identity (**Extended Data Fig. 4a; SI Table 2**). Ingenuity pathway analyses of DEGs between young and old populations revealed repression of extracellular matrix genes, including many collagens, and activation of several biosynthetic pathways in old AEC-like cells (**Extended Data Fig. 4b, c; SI Table 3**). In contrast, old SEC-like cells showed a broad downregulation of signaling pathways, decreased expression of genes involved in endocytosis, and activation of several cell death pathways. These molecular alterations underpin the fragilized and dysfunctional nature of old SECs, and demonstrate an overall loss of vascular integrity with age that also affects the arteriolar network.

Finally, we complemented our targeted single-cell niche profiling of the old niche by performing unbiased droplet-based scRNAseq on unfractionated Ter119^−^/CD45^−^ endosteal and marrow fractions isolated from young and old mice. UMAP representation combining both location confirmed the severe degradation of EC and endosteal MSC-S/OPr compartments associated with induction of cell death pathways and osteo-chondrocytic differentiation block, as well as the major expansion of inflammatory marrow MSC-L in old mice (**Fig. 2c; Extended Data Fig. 5a**). Interestingly, we did not detect activation of senescence-related gene expression programs in any old mesenchymal populations, which was surprising since the changes in old BM fluids displayed a clear overlap with known senescence-associated secretory pathway (SASP) cytokines^32^ (**Extended Data Fig. 5b**). Isolated old endosteal MSC-S and OBC were also negative for senescence-associated β-galactosidase (SA-β-gal) staining (**Extended Data Fig. 5c**). Finally, a qRT-PCR survey of various hematopoietic cells including macrophages did not uncover significant changes in *Il1a*, *Il1b*, or *Tnf* mRNA expression with age, in contrast to unfractionated CD45^−^/Ter-119^−^ endosteal stroma that showed elevated *Il1b* and *Tnf* expression (**Extended Data Fig. 5d**). Plate-based scRNAseq analyses further confirmed that discrete stromal populations including degraded endosteal OPr and AEC, and expanded marrow iMSC-L, contributed to the inflamed milieu in the old BM cavity (**Extended Data Fig. 5e**). Taken together, these molecular results confirm that the remodeling of the old niche contribute to the establishment of an inflamed BM milieu.

## Inflammatory remodeling of the old blood system

To address the consequence of niche inflammation on the hematopoietic system, we first compared steady state hematopoiesis in young and old mice (**Extended Data Fig. 6a-d**). Old mice displayed elevated levels of myeloid and lymphoid cells, relatively unchanged red blood cell (RBC) levels and increased platelet (Pt) counts in peripheral blood that mirror the accumulation of mature Mks and Mk progenitors (MkP) in the BM (**Extended Data Fig. 6e-g**). In contrast, the numbers of mature myeloid cells and committed myeloid progenitors in the BM remained fairly similar between young and old mice, while the number of common lymphoid progenitors (CLP) and BM resident B cells significantly decreased with age. Hematopoietic aging has long been characterized by an expansion of the Lin^−^/c-Kit^+^/Sca-1^+^ (LSK) compartment^1,2^, which contains HSCs and all of the MPP subsets^5^. This reflected a massive expansion of CD34^−^/CD41^+^ Mk-biased HSCs that also expressed other Mk-lineage markers like P-selectin and von Willebrand Factor (vWF)^28,33^, as well as increased production of myeloid-biased MPP2 and MPP3, and decreased numbers of lymphoid-biased MPP4^34^ (**Fig. 3a, b**). These changes confirm alterations in old HSC differentiation potential at steady state.

**Figure 3.**
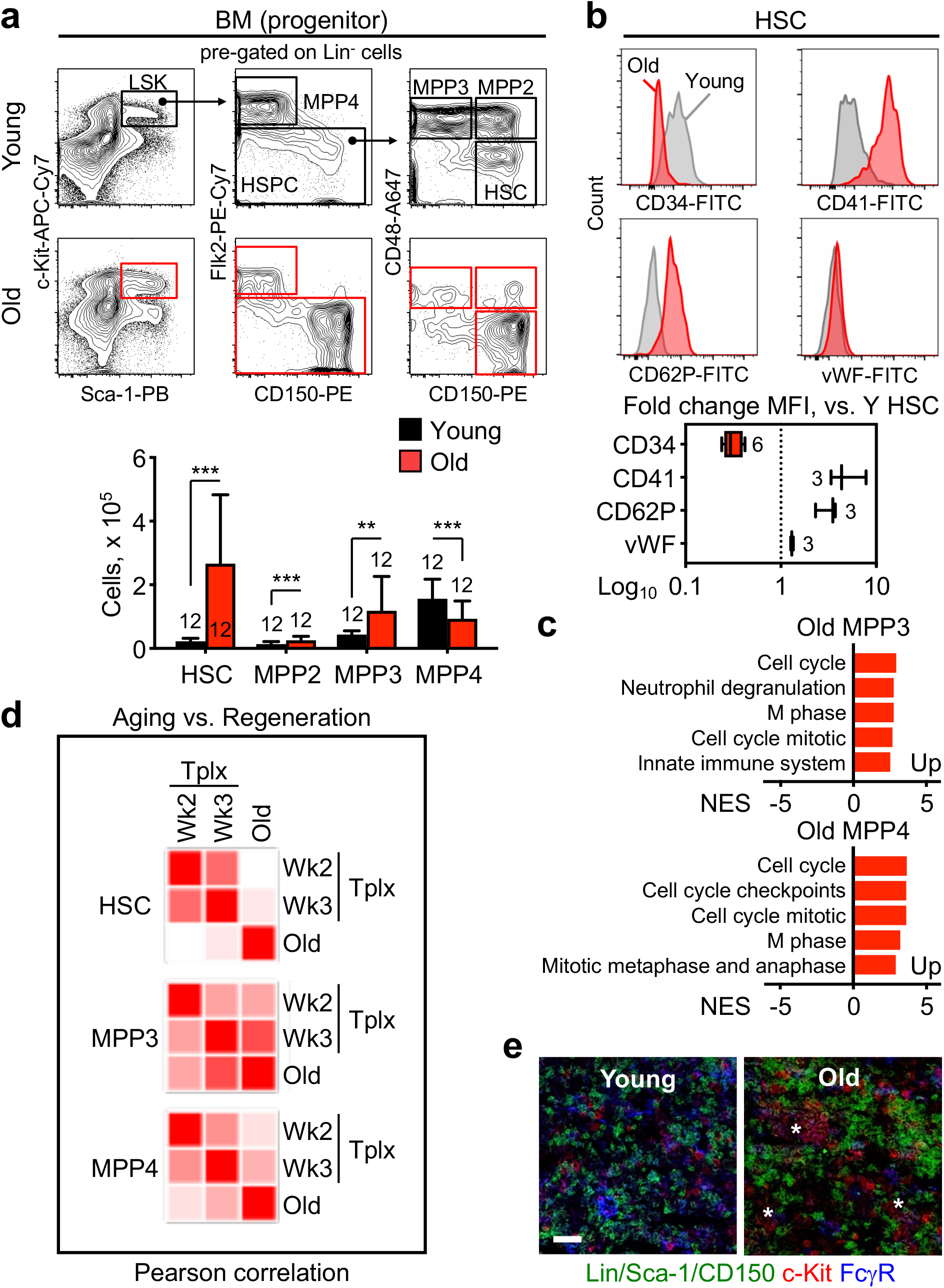
Activation of emergency myelopoiesis pathways in the old blood system. **a,** Representative flow cytometry plots and quantification of HSCs and MPP populations in young and old mice. **b**, Representative staining and quantification of CD34, CD41, CD62P and vWF levels on young and old HSCs. **c**, GSEA analyses of gene expression microarray data from young and old HSCs, MPP3 and MPP4 (3 biological replicates per population). **d**, Pearson correlation of Fluidigm gene expression data comparing old and regenerative HSCs, MPP3 and MPP4. Regenerating populations were isolated 2 and 3 weeks following young HSC transplantation (Tplx)^5^. **e,** Representative image of GMP immunofluorescence staining in young and old BM. Stars indicate GMP patches. Scale bar, 60 μm. Data are means ± S.D; **p ≤ 0.01, ***p ≤ 0.001.

To understand the consequences of those changes for downstream progenitor function, we next investigated old MPP populations. *In vitro*, old MPP3 showed reduced clonogenic potential in methylcellulose and accelerated myeloid differentiation in liquid culture (**Extended Data Fig. 7a, b**). In contrast, old MPP4 had elevated myeloid colony-forming activity and, similarly to MPP3 and HSCs, delayed B cell differentiation potential in OP9/IL-7 co-cultures (**Extended Data Fig. 7c**). However, none of the old MPP populations exhibited delayed division kinetics associated with replication stress^35^ or dampened apoptotic responses^36^ characteristic of old HSCs (**Extended Data Fig. 7d, e**). Transplantation and short-term lineage-tracking in sub-lethally irradiated recipients confirmed the reduced reconstitution activity from old MPP3, and highlighted a strong myeloid bias from all old populations including MPP4 (**Extended Data Fig. 8a**). In addition, terminal analyses at 2 and 3 weeks post-transplantation revealed no difference in the ability of young and old HSCs to produce donor-derived HSCs, MPP3 and MPP4 in young recipients (**Extended Data Fig. 8b**), suggesting that they are both responsive to hematopoietic stress. Altogether, these results indicate that old MPPs, while clearly altered in their lineage differentiation potential, do not share most cell-intrinsic hallmarks of HSC aging.

To gain insights into the molecular mechanisms responsible for the altered functionality of old MPPs, we took advantage of our previously performed microarray gene expression analyses^5^ to include various old progenitor populations. Targeted GSEA demonstrated upregulation of myeloid genes with a concomitant decrease in CLP genes in old MPP3, and enrichment for preGM myeloid progenitor genes in old MPP3 and MPP4 (**Extended Data Fig. 8c**). Untargeted analyses further established the unique signature of old MPPs, with old MPP4 upregulating cell cycle and DNA repair genes, and old MPP3 also expressing genes specific to the innate immune response (**Fig. 3c; SI Table 4**). Complementary Fluidigm-based qRT-PCR analyses comparing aging and previously published regenerative signatures^5^ highlighted the correlation between old and 3 week post-transplantation MPP3 and MPP4, and to a lower extent HSCs, further suggesting that aging constitutively triggers normally transient myeloid regeneration programs (**Fig. 3d**). Finally, we observed the presence of GMP patches in the old marrow cavity (**Fig. 3e**), which are never found in young mice absent a regenerative challenge^37^. Together, these results demonstrate a profound hematopoietic remodeling with constitutive activation of emergency myelopoiesis pathways from old HSCs with increased production of MPP3, myeloid-reprogramming of MPP4, and GMP cluster formation likely caused by persistent exposure to niche inflammation.

## Impaired hematopoietic recovery from stress

To understand how the old blood system responds to hematopoietic stress, we injected young and old mice with 5-fluoruracil (5FU). Old mice were exquisitely sensitive to myeloablation with ~ 50% mortality likely caused by massive thrombosis and anemia (**Extended Data Fig. 9a, b**). Blood analyses revealed a severely altered regenerative response with age, with a dramatic rebound and overproduction of mature myeloid cells, B cells and Pts associated with a severe and persistent loss of RBCs (**Extended Data Fig. 9b-d**). BM analyses confirmed increased myeloid cell production in response to 5FU challenge in old mice, but uncovered decreased B lymphopoiesis suggesting that a purging from the BM rather than increased production contributed to their elevated circulating levels. In addition, we observed an amplified HSC response associated with increased and significantly delayed MPP and downstream progenitor amplification, delayed GMP cluster differentiation, and impaired erythroid progenitor production in 5FU-treated old mice (**Extended Data Fig. 9e, f**). Quantification of the local cytokine milieu indicated an exacerbation of 5FU-induced niche inflammation, with elevated IL-1α, IL-1β and MIP-1α levels found in the old marrow cavity (**Extended Data Fig. 9f**). Altogether, these results suggest that by constitutively activating emergency myelopoiesis from old HSCs, the old niche hinders hematopoietic regeneration and exacerbates blood aging features hence furthering myeloid bias at the expense of other lineages.

## Improved recovery upon blocking IL-1 signaling

To demonstrate the functional impact of age-related niche inflammation, we focused on IL-1, a well-known regulator of myeloid regeneration^6^. Chronic IL-1β exposure in young mice is known to mimic features of blood aging^38^, with myeloid cell expansion and decreased lymphopoiesis associated with anemia and thrombosis (**Extended Data Fig. 10a**). Strikingly, chronic IL-1β exposure also phenocopies several effects of aging on niche cells, with expansion of a dysmorphic and leaky sinusoidal network in the central marrow associated with SEC loss upon cell isolation, and decreased numbers of endosteal MSC-S and OBC (**Fig. 4a-c**). These results indicate that elevated BM IL-1 levels observed with age directly contribute to the functional deterioration of both the niche and blood system.

**Figure 4.**
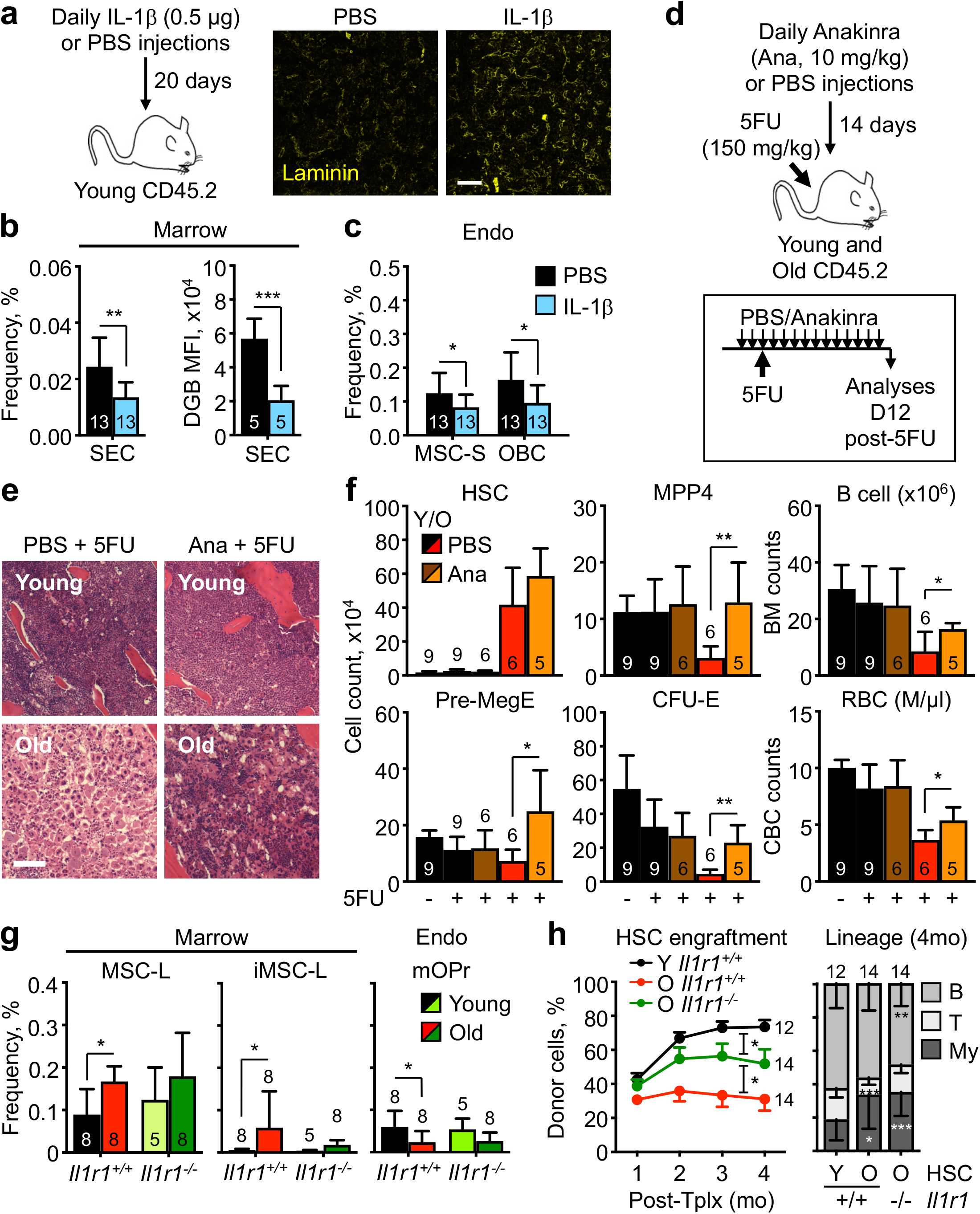
IL-1 signaling drives the aging of both niche and blood system. **a-c,** Pro-aging effects of chronic IL-1β exposure in young mice with: (a) experimental scheme and representative staining of the BM vasculature (scale bar, 100 μm); (b) changes in SEC frequency and dragon green beads (DGB) retention; and (c) changes in endosteal mesenchymal populations. **e-f,** Rejuvenating effects of IL-1 signaling blockade with anakinra (Ana) during 5FU-mediated regeneration in old mice with: (d) experimental scheme; (e) representative H&E staining of sternum (scale bar, 100 μm); and (f) quantification of the indicated BM and blood populations. Results are from 3 independent cohorts with a total of 15 young and 11 old mice treated once with 5FU and injected with either PBS or Anakinra. **g-h,** Improved aging features in old *Il1r1^−/−^* mice with: (g) changes in central marrow and endosteal mesenchymal population frequencies; and (h) engraftment over time (left, mean ± S.E.M.) and lineage reconstitution (right) at 4 months (4 mo) post-transplantation (Tplx) of the indicated HSC populations (250 HSCs per mouse). Results are from 3 independent cohorts of young and old *Il1r1^−/−^* mice and age-matched *Il1r^+/+^* controls. Data are means ± S.D except when indicated; *p ≤ 0.05, **p ≤ 0.01, ***p ≤ 0.001.

To determine whether targeting IL-1 signaling could revert aging features, we first administered the human IL-1 receptor antagonist Anakinra (Kineret) intraperitoneally for 14 days in young and old mice (**Extended Data Fig. 10b**). However, short-term steady state inhibition did not significantly alter the composition of the HSC/MPP pool, nor the defective engraftment and lineage bias of old HSCs (**Extended Data Fig. 10c, d**). We next investigated whether blocking IL-1 signaling could improve hematopoietic regeneration (**Fig. 4d**). Strikingly, daily treatment with Anakinra before and after 5FU injection ameliorated BM regeneration in old mice, alleviating anemia and improving B lymphopoiesis (**Fig. 4e, f; Extended Data Fig. 10e**). Effective B cell and RBC recovery was accompanied by increased MPP4 and CFU-E production, respectively (**Fig. 4f**). BM fluid analyses also confirmed a specific reduction in IL-1α and IL-1β levels in Anakinra-treated 5FU-challenged old mice, demonstrating the efficacy of the treatment (**Extended Data Fig. 10f**). To test the necessity of IL1 signaling in driving aging phenotypes, we finally analyzed young and old *Il1r1^−/−^* mice together with age-matched *Il1r1^+/+^* controls. Old *Il1r1^−/−^* mice showed reduced features of blood aging, with limited myeloid cell expansion, B cell loss and anemia in the blood, and attenuated MPP3 expansion and myeloid cell production in the BM, despite persistence of an expanded HSC compartment (**Extended Data Fig. 10g, h**). Old *Il1r1^−/−^* mice also displayed features of a more youthful niche, with limited loss of endosteal mOPr and reduced expansion of marrow MSC-L and iMSC-L (**Fig. 4g**). Moreover, old *Il1r1^−/−^* HSCs exhibited increased fitness in transplantation assays with significantly improved overall chimerism (**Fig. 4h**). Together, these results demonstrate a direct role for IL-1 in driving the aging of both the niche and blood system, and indicate that blocking chronic IL-1 signaling offers a therapeutic route to revert age-related lineage biases and improve hematopoietic regeneration in chemotherapeutic and transplantation settings.

## Discussion

Chronic inflammation is a hallmark of organismal aging but its consequences for tissue function often remain unclear^9^. Here, we demonstrate that BM aging is defined by the remodeling of the microenvironment with increased production of pro-inflammatory cytokines in large part by dysfunctional niche cells and activation of immune response programs in both hematopoietic and stromal cells. Inflammaging directly contributes to the loss of endosteal mesenchymal populations, impaired osteogenesis, and disorganization of sinusoidal blood vessels with increased vascular leakiness. These changes, along with an expansion of inflammatory perisinusoidal MSC-L, lead to a self-reinforcing cycle of damage that drives lineage biases and regenerative response defects from the aged blood system. In particular, localized inflammation drives the constitutive activation of emergency myelopoiesis pathways from old HSCs and MPPs, reinforcing myeloid cell production at the expense of lymphoid and erythroid commitment. This in turn blunts regenerative responses that rely on acute activation of these pathways, leading to an exacerbated phenotype in stress conditions. Our results provide a novel understanding of blood aging based on crosstalk between the inflamed niche and the inflamed hematopoietic system leading to degraded blood production both at steady state and during regeneration.

Our results identify IL-1 as a major and targetable driver of age-related niche and blood system deterioration. In fact, elevated IL-1β levels and activation of inflammasomes have been demonstrated in human studies and correlated with increased age-related mortality^39^. Here, we show that blocking IL-1 signaling can revert many differentiation biases from old HSCs and improve blood recovery from stress, while also attenuating features of aging in the old niche. Our results offer a potential therapeutic application of IL-1 inhibitors to improve blood production in the elderly, especially when hematopoietic regeneration is needed following chemotherapy or other immunosuppressive treatments. Future experiments will establish the precise role of other inflammatory mediators increased with age in the marrow cavity, and determine whether combined interventions can further delay BM niche aging and recover youthful blood production.

## Supplementary information

See accompanying document.

## Acknowledgments

We thank S. Villeda (UCSF) for providing some old C57Bl/6 mice, D. Reynaud (UCSF) for initial analyses of old mice, A. Valencia (UCSF) for technical assistance with various experiments, S. Kinston (Cambridge) for scRNAseq library preparations, A. Li (Bone Imaging Research Core, UCSF) for μCT analyses, M. Lee (UCSF) and M. Kissner (CUIMC) for management of our Flow Cytometry Core facilities, and all members of the Passegué laboratory for critical insights and suggestions. E.V.V. was supported by a Rubicon Grant from The Netherlands Organization for Scientific Research, a Stem Cell Grant from BD Biosciences, and a NYSTEM training grant, P.V.D by NIH F31 HL151140, E.M.P. by NIH F32 HL106989 and K01 DK09831, S.T.B. by a CIRM postdoctoral fellowship, and T.T.H. by an AHA and Hillblom Center for the Biology of Aging predoctoral fellowships. F.J.C-N., X.W. and B.G. were supported by grants from the Wellcome (206328/Z/17/Z), CRUK (C1163/A21762) and core funding by the Wellcome to the Cambridge Stem Cell Institute. This work was funded by NIH R01CA184014, NIH R35HL135763, Glenn Foundation Research Award and LLS Scholar Award to E.P., and supported in part through the NIH/NCI Cancer Center Support Grant P30CA013696 to CUIMC. For the purpose of Open Access, the author has applied a CC BY public copyright license to any Author Accepted Manuscript version arising from this submission.

## Author Contributions

E.V.V. contributed to most of the experiments with help from A.H. for BMF isolation and some immunophenotyping, S.Y.Z. for GMP cluster analyses and ELISA assays, A.F.S. for immunofluorescence and whole-mount staining, S.T.B for initial 5FU analyses, and T.T.H. for cell harvest and technical assistance. C.A.M and P.V.D contributed to some stromal investigations and analyzed *Il1r1*-deficient mice. F.J.C-N., X.W. and B.G. prepared and analyzed the scRNA-seq samples, and E.M.P. prepared the microarray samples. C.A.M. re-analyzed all the generated data and performed pathway analyses. E.V.V and E.P. designed the experiments and together with C.A.M. wrote and edited the manuscript.

## Competing interests

The authors declare no competing financial interests.

## Additional information

Supplementary information is available for this paper. Correspondence and requests for materials should be addressed to E.P.

## Methods

### Mice

Young and old wild-type C57BL/6-CD45.2 (B6) and wild-type C57BL/6-CD45.1 (BoyJ) mice of both sexes were bred and aged in house either at UCSF or CUIMC. Some old B6 mice were also obtained from the National Institute on Aging (NIA) aged rodent colonies and from collaborators. β-actin–GFP C57BL/6-CD45.2 transgenic mice^5^ and *Il1r1^−/−^* C57BL/6-CD45.2 mice^38^ were previously described. At the time of analyses, young mice were 6-12 weeks of age, middle-age mice were 13 months old, and old mice were all ≥ 22 months of age, most of them ~ 24 month of age and some of them 26-31 months of age for specific experiments. Young BoyJ recipient mice for HSC and MPP primary and secondary transplantation assays were 8-12 weeks of age at time of irradiation. No specific randomization or blinding protocol was used with respect to the identity of experimental animals, and both male and female animals were used indiscriminately in all experiments. All mice were maintained in mouse facilities at UCSF or CUIMC in accordance with IACUC protocols approved at each institution.

### *In vivo* assays

For quantitative Dragon Green Bead (DGB; Bangs Laboratories, FSDG001) analyses, mice were injected retro-orbitally with 2.5 μl/g DGB solution under isoflurance anaesthesia 10 min prior to euthanasia by CO2 asphyxation and immediately perfused with 20 ml PBS by cardiac puncture before bone harvest. For 5-fluorouracil (5FU; Sigma-Aldrich) treatment, mice were injected intraperitoneally with 150 mg/kg 5FU or vehicle (PBS) and analyzed for blood and BM parameters. For chronic IL-1 treatment, mice were injected intraperitoneally with 0.5 μg IL-1β (Peprotech, 211-11B) in 100 μl PBS/0.2% BSA or vehicle (PBS/0.2% BSA) daily for 20 days and analyzed for blood and BM parameters. For Anakinra treatment, mice were injected intraperitoneally with 10 mg/kg Anakinra (Swedish Orphan Biovitrum AB, 666-58234-07) or vehicle (PBS) daily for 14 days, with or without 5FU injection on the third day, and analyzed for blood and BM parameters. For transplantation experiments, CD45.1 recipient mice were exposed to 9 Gy (sub-lethal) or 11 Gy (lethal) irradiation dose delivered in split doses 3 hours apart using either a 137Cs source (J. L. Shepherd) or an X-ray irradiator (MultiRad225, Precision X-Ray Irradiation), and purified HSCs and MPPs were delivered via retro-orbital injection. For transplantation into sub-lethally-irradiated recipients, mice were injected with 2,000-5,000 purified CD45.2 HSCs or MPPs. For transplantation in lethally-irradiated recipients, mice were injected with 250 CD45.2 HSCs delivered together with 300,000 Sca-1-depleted CD45.1 helper BM cells. Recipient mice were administered polymyxin/neomycin-containing water for 4 weeks following the procedure to prevent opportunistic infection, and analyzed for blood and BM parameters. Peripheral blood was collected under isoflurane anesthesia via retro-orbital bleeding and dispensed into EDTA-coated tubes (Becton Dickinson) for complete blood count (CBC) analyses using a Genesis (Oxford Science) hematology system. BM analyses were terminal analyses at the time of tissue harvest and BM cellularity was determined using a ViCell automated cell counter (Beckman-Coulter).

### Flow cytometry of hematopoietic cells

BM hematopoietic stem, progenitor and mature cell populations were analyzed and/or purified as previously described^5^. In brief, BM cells were obtained by crushing leg, arm, and pelvic bones (with sternum and spines for some experiments) in staining media composed of Hanks’ buffered saline solution (HBSS) containing 2% heat-inactivated FBS (Cellgro B003L52). Red blood cells were removed by lysis with ACK (150 mM NH4Cl/10 mM KHCO3) buffer, and single-cell suspensions of BM cells were purified on a Ficoll gradient (Histopaque 1119, Sigma-Aldrich). For HSC and progenitor isolation, BM cells were pre-enriched for c-Kit^+^ cells using c-Kit microbeads (Miltenyi Biotec, 130-091-224) and an AutoMACS cell separator (Miltenyi Biotec). Unfractionated or c-Kit-enriched BM cells were then incubated with purified rat anti-mouse lineage antibodies (CD3, BioLegend, 100202; CD4, eBioscience, 16-0041-82; CD5, BioLegend, 100602; CD8, BioLegend, 100702; CD11b, BioLegend, 101202; B220, BioLegend, 103202; Gr1, eBioscience, 14-5931-85; Ter119, BioLegend, 116202) followed by goat anti-rat-PE-Cy5 (Invitrogen, A10691) and subsequently blocked with purified rat IgG (Sigma-Aldrich). Cells were then stained with c-Kit-APC-Cy7 (BioLegend, 105826), Sca-1-PB (BioLegend, 108120) or Sca-1-BV421 (BioLegend, 108128), CD150-PE (BioLegend, 115904) or CD150-BV650 (BioLegend, 115931), CD48-A647 (BioLegend, 115904) or CD48-A700 (BioLegend, 103426), and Flk2-biotin (eBioscience, 13-1351-85) or Flk2-PE (eBioscience, 12-1351-82) followed by SA-PE-Cy7 (eBioscience, 25-4317-82) for HSC/MPP staining, or together with CD34-FITC (eBioscience, 11-0341-85) or CD34-biotin (BioLegend, 119304) and FcγR-PerCP-Cy5.5 (eBioscience, 46-0161-82) or FcγR-PE-Cy7 (BioLegend, 101317) followed by SA-BV605 (BioLegend, 405229) for combined HSC/MPP/myeloid progenitor staining. For quantification of HSC surface marker expression, CD41-FITC (eBioscience, 11-0411-82), CD62P-FITC (BD, 561923) and vWF-FITC (EMFRET, P150-1) were also used. For extended myeloerythroid progenitor staining, unfractionated BM cells were incubated with Lin/PE-Cy5 and then CD41-PE (BD, 558040) or CD41-BV510 (BioLegend, 133923), FcγR-PerCP-Cy5.5 or FcγR-PE-Cy7, CD150-APC (BioLegend, 115910) or CD150-BV650, and CD105-A488 (BioLegend, 120406) or CD105-BV605 (BD, 740425). For CLP staining, unfractionated BM cells were incubated with Lin/PE-Cy5, then cKit-APC-Cy7, Sca-1-PB, Flk2-biotin, and IL7R-PE (eBioscience, 12-1271-82) followed by SA-PE-Cy7. For mature cell analyses, depending on the experiments, BM cells were stained with Mac-1-PE-Cy7 (eBioscience, 25-0112-82), Gr-1-e450 (eBioscience, 57-5931-82), B220-APC-Cy7 (eBioscience, 47-0452-82), and CD3-FITC (BioLegend, 100306). For myeloid cell isolation, BM cells were stained with CD3-PE-Cy5 (eBioscience, 15-0031-63), B220-PE-Cy5 (eBioscience, 15-0452-82), NK1.1-PE-Cy5 (BioLegend, 108716), CD11b-FITC (eBioscience, 11-0112-82), Ly-6C-APC-Cy7 (BioLegend, 128025) and Ly-6G-A700 (BioLegend, 127622). For lymphoid cell isolation, BM cells were stained with B220-APC-Cy7, CD19-PE (eBioscience, 12-0193-82), TCRβ-PE-Cy7 (Invitrogen, 25-5961-80), CD4-PE-Cy5 (eBioscience, 15-0041-82), and CD8-A700 (Pharmigen, 557959). For *in vitro* differentiation assays, cultured cells were stained with c-Kit-APC-Cy7, Sca-1-PB, Mac-1-PE-Cy7 and FcγR-PE (eBioscience, 12-0161-83) for myeloid differentiation, or Mac-1-APC (eBioscience, 17-0112-82) and CD19-PB (BioLegend, 115523) for lymphoid differentiation on OP9 cells. For HSC chimerism analyses, HSCs were stained with Lin/PE-Cy5, c-Kit-APC-Cy7, Sca-1-BV421, CD150-BV650, CD48-A700, and Flk2-bio followed by SA-BV605, together with CD45.2-FITC (eBioscience, 11-0454-85) and CD45.1-PE (eBioscience, 25-0453-83). For peripheral blood chimerism analyses, cells were stained with Mac-1-PE-Cy7, Gr-1-e450, B220-APC-Cy7, CD3-APC (eBioscience, 17-0032-82), and Ter-119-PE-Cy5 (eBioscience, 15-5921-83) together with CD45.2-FITC and CD45.1-PE. Before isolation or analyses, stained cells were resuspended in staining media containing 1 μg/ml propidium iodide (PI) for dead cell exclusion. HSC and progenitor cell isolations were performed on a Becton Dickinson (BD) FACS Aria II (UCSF) or FACS Aria II SORP (CUIMC) using double sorting for purity. Mature cell isolations were performed on a FACS Aria II SORP using single sorting on purity mode. Cell analyses were performed either on the same FACS ARIA, or on a BD LSR II (UCSF), BD Celesta (CUIMC) or Bio-Rad ZE5 (CUIMC) cell analyzer. For all experiments, HSCs were identified as Lin^−^/Sca-1^+^/c-Kit^+^/Flk2^−^/CD48^−^/CD150^+^ BM cells, MPP3 as Lin^−^/Sca-1^+^/c-Kit^+^/Flk2^−^/CD48^+^/CD150^+^ BM cells, MPP4 as Lin^−^/Sca-1^+^/c-Kit^+^/Flk2^+^ BM cells, and, except when indicated, granulocytes (Gr) cells as Mac-1^+^/Gr-1^+^ BM cells, B cells as B220^+^ BM cells and T cells as CD3^+^ BM cells.

### Flow cytometry of niche cells

Endosteal stromal cells were isolated as previously described^30^. In brief, leg, arm, and pelvic bones were gently crushed and washed with HBSS until white. Stromal cells were released by treatment with 3 mg/ml Type I Collagenase (Worthington) in 2 ml HBSS for 1 hr at 37°C with shaking at 110 rpm. Stromal cells were washed with HBSS + 4% FBS and filtered through 45 μm mesh into a polypropylene tube. Isolation of central marrow stromal cells was adapted from a published protocol^40^ using individual femurs with the femoral head cut off and kneecap removed to expose the growth plate. Intact BM plugs were carefully flushed with HBSS into polypropylene tubes by inserting 1 ml syringes with 22G needles into the growth plate. HBSS was discarded and stromal cells were released by digestion with 3 mg/ml Type I Collagenase (Worthington) in 1 ml HBSS for 10 min twice at 37°C with shaking at 110 rpm. After the second incubation the plugs were resuspended by pipetting and filtered through 45 μm mesh into a new tube. Both endosteal and central marrow stromal cells were stained with Ter119-PE-Cy5, CD45-APC-Cy7 (BD, 557659), CD31-PE (BD, 553373), Sca-1-A700 (eBioscience, 56-5981-82), CD105-BV786 (BD, 564746), CD51-BV421 (BD, 740062), PDGFRα-PE-Cy7 (eBiosciences, 25-1401-82) and LepR-biotin (R&D Systems, BAF497) followed by SA-APC (eBiosciences, 17-4317-82), and resuspended in HBSS with 4% FBS and PI. Both cell isolations and analyses were performed on a Becton Dickinson (BD) FACS Aria II (UCSF) or FACS Aria II SORP (CUIMC) using single sorting on purity mode. For all experiments, MSC-S were identified as Ter119^−^/CD45^−^/CD31^−^/Sca-1^+^/CD51^+^, OBC as Ter119^−^/CD45^−^/CD31^−^/Sca-1^−^/CD51^+^, mOPr as Ter119^−^/CD45^−^/CD31^−^/Sca-1^−^/CD51^+^/PDGFRα^+^/LepR^−^, OPr as Ter119^−^/CD45^−^/CD31^−^/Sca-1^−^/CD51^+^/PDGFRα^−^/LepR^−^ and AEC as Ter119^−^/CD45^−^/CD31^+^/Sca-1^hi^/CD105^lo^ endosteal cells, while MSC-L were identified as Ter119^−^/CD45^−^/CD31^−^/Sca-1^−^/CD51^+^/PDGFRα^+^/LepR^+^ and SEC as Ter119^−^/CD45^−^/CD31^+^/Sca-1^lo^/CD105^hi^ central marrow cells.

### *In vitro* assays

All cultures were performed at 37 °C in a 5% CO2 water jacket incubator (ThermoFisher). For methylcellulose colony assays, single cells were directly sorted into individual wells of a flat-bottom 96-well plate (Fisher Scientific, 353072) containing 100 μl methylcellulose (Stem Cell Technologies, M3231) supplemented with penicillin (50 U/ml)/streptomycin (50 μg/ml), 0.1 mM non-essential amino acids (Fisher Scientific, 11-140-050), 1 mM sodium pyruvate (Fisher Scientific, 11-360-070), 2 mM L-glutamine (Fisher Scientific 35-050-061), 50 μM 2-mercaptoethanol (Sigma, M7522) and the following cytokines (all from PeproTech): IL-3 (10 ng/ml), GM-CSF (10 ng/ml), SCF (25 ng/ml), IL-11 (25 ng/ml), Flt-3L (25 ng/ml), TPO (25 ng/ml) and EPO (4 U/ml). Colonies were visually scored after 7 days of culture. For myeloid differentiation assays, 1,000 cells were directly sorted into individual wells of a 96-well plate containing Iscove’s modified Dulbecco’s media (IMDM) medium (Invitrogen) suplemented with 5% FBS (StemCell Technology, 06200) and the same reagents and cytokine cocktail than the methylcellulose assays (full cytokine medium). Cells were analyzed by flow cytometry after different culture periods. For cleaved caspase 3/7 (CC3/7) assays, 400-600 cells were directly sorted in triplicate into 40 μl full cytokine medium in a 384-well plate, and incubated for 24 hr before adding 40 μl of Caspase-Glo 3/7 (Promega) to each well. Plates were then shaken for 30 s at 500 rpm, incubated for 45 min at RT and read on a luminometer (Synergy2, BioTek) to obtain relative units (RU). OP9 stromal cells (ATCC, CRL-2749) were maintained in Minimum Essential Medium Eagle alpha modification (αMEM) medium (Invitrogen) supplemented with 10% FBS (Cellgro, B003L52), L-glutamine (2 mM), and penicillin (50 U/ml)/streptomycin (50 μg/ml), and split at 1:4 every 3-4 days as needed to avoid over-confluence. For lymphoid differentiation assays, 500 cells were directly sorted into individual wells of a 24-well plate containing 10,000 OP9 stromal cells in OptiMEM medium (Invitrogen) supplemented with 5% FBS (StemCell Technology), L-glutamine (2 mM), penicillin (50 U/ml)/streptomycin (50 μg/ml), 2-mercaptoethanol (50 μM), SCF (10 ng/ml), Flt3L (10 ng/ml), and IL-7 (5 ng/ml). Following sequential withdrawal of Flt3L and SCF upon 2-day intervals, cultures were maintained in IL-7 and analyzed by flow cytometry at various intervals. For CFSE analyses, 1,000-1,500 cells were directly sorted into individual wells of a 96-well plate containing staining media, washed with PBS, resuspended in 100 μl PBS containing 5 μM CFSE (Molecular Probes, C-1157), incubated for 5 minutes at RT, quenched with FBS, incubated for 1 min at RT, washed twice with staining media, resuspended in 200 μl full cytokine medium, incubated for 72 hr and analyzed by flow cytometry. Stromal cells were grown in αMEM supplemented with 10% FBS (CellGro), penicillin (50 U/ml)/streptomycin (50 μg/ml), and 2-mercaptoethanol (50 μM).

For fibroblastic colony-forming unit (CFU-F) assays, endosteal MSC-S (15-300 cells) or OBC (170-300 cells) were sorted directly in 6-well plates containing 1.5 ml medium/well and cultured for 11 days with medium exchange every 2-3 days, before staining with Giemsa-Wright to score colonies of 25 or more cells. In contrast, marrow MSC-L (100-300 cells) were sorted directly into 6-well plates containing 1.5 ml of Dulbecco’s Modified Eagle Medium (Gibco) supplemented with 20% FBS (CellGro), penicillin (50 U/ml)/streptomycin (50 μg/ml), 2-mercaptoethanol (50 μM), and 10 μM ROCK inhibitor (TOCRIS, Y-27632) and cultured in hypoxic conditions (5% O_2_) for 8 days with medium exchange every 2-3 days, before staining with Giemsa-Wright to score colonies of 25 or more cells. For co-culture assays, 25 β-actin-GFP MSC-S cells were directly sorted into 96-well plates containing 100 μl medium/well, then 1×10^5^ unfractionated BM cells in 100 μl medium were added and co-cultures were incubated for 10 days without medium replacement. On day 10, medium was aspirated and adherent cells were washed twice with PBS, liberated with 50 μl 0.25% trypsin/EDTA (Thermo, 25200056), and counted by flow cytometry using 25 μl Absolute Counting Beads (Life Technologies, C36950). For SA-β-Gal (Cell Signaling, 9860S) staining, MSC-S and OBC were sorted directly onto poly-L-lysine coated slides (Sigma, P0425) and stained according to manufacturer’s protocol.

### Bone analyses

Bones for hematoxylin/eosin (H&E) staining were processed by the Mouse Pathology Core Facility at UCSF or the Molecular Pathology Core Facility at CUIMC. Bones for MicroCT analyses were fixed in 10% neutral buffered formalin for 24 hours, then held in 70% ethanol at 4°C until further processing. CT scans were performed on a vivaCT40 MicroCT (Scanco; 55 kV x-ray energy, 10.0 μm voxels, 500 ms integration times) and bone densitometry analyses were performed as previously described^33^. Trabecular tissue volume (TV), mineralized bone volume (BV), and trabecular bone parameters were determined on 100 slices, starting 500 μm below the bottom edge of the growth plate using segmentation values of 0.5/2/350 which corresponds to 650 mg hydroxyapatite/cm^3^.

### Immunofluorescence staining

For whole mount imaging, mice were injected retro-orbitally under isoflurane anesthesia with 100 μl of CD31-A647 (BioLegend, 102416) and VE-Cadherin-A647 (BioLegend, 138006) antibody cocktail in PBS. Injected mice were euthanized 10 min later, perfused with 3 ml PBS and fixed by perfusion of 10 ml 4% paraformaldehyde (PFA) in PBS. Femurs were harvested, fixed in 4% PFA for 2 hr on ice, washed 3x with PBS for 10 min each, and then cleared by successive dehydration with sucrose (15% and then 30%) for either 1 hr or overnight at each step. Bones were then snap-frozen in a 100% ethanol/dry ice slurry, and kept at −80 °C until sectioning. One side of the frozen femura was cryosectioned with a tungsten blade (Leica, CM3050 S) until the BM was exposed, then transferred to a 1.5mL Eppendorff tube, washed 3x with PBS at RT and blocked with 20% goat-serum in PBS/0.1% Tween-20 (PBST) overnight at 4°C. Bones were incubated with rabbit anti-mouse laminin (Sigma, L9393) in 10% goat-serum in PBST for 3 days at 4°C, then washed 3x with PBST and incubated with donkey goat anti-rabbit A594 (Thermo Fisher, A32740) for 2 days at 4°C. Finally, bones were washed 3x with PBST for 10 min each, mounted on a cover-slip with silicone glue, kept wet with PBS at all times, and imaged on an SP8 inverted confocal microscope (Leica) with 20x objective. For BM section imaging, mouse femurs were embedded in OCT, snap-frozen in a 100% ethanol/dry ice slurry, and kept at −80 °C until sectioning. Thin 7 μm section were obtained upon cryosection using the CryoJane tape transfer system (Leica, 39475205) and a tungsten blade. Sections were dried for 2-4 hr at room temperature and then frozen at −80 °C until stained. Before staining, sections were fixed with 100% acetone for 10 min at −20 °C, dried for 5 min at RT, blocked for 90 min with 10% goat-serum (Gibco) in PBS and washes 3x with PBS. For LepR staining, sections were first stained with LepR-biotin (R&D systems, AF497) and rabbit anti-mouse laminin, and then with SA-A555 (ThermoFisher, S21381) and goat anti-rabbit A594 (Thermo Fisher, A32740) followed by blocking with rat IgG for 10 min at RT. Slides were then stained with A488-conjugated lineage markers Ter119 (BioLegend, 116215), Mac-1 (BioLegend, 101217), Gr-1 (BioLegend, 108417), B220 (BioLegend, 103225), CD41 (Thermo Fisher, 11-0411-82) and CD3 (BioLegend, 100210). For ALCAM staining, sections were first stained with anti-ALCAM-Bio (R&D systems, BAF1172) followed by SA-A555 blocked with rat IgG and then stained with A488-conjugated lineage markers. For laminin staining, sections were stained with rabbit anti-mouse laminin followed by goat anti-rabbit-A647 and, eventually, DGB fluorescence was detected in the A488 channel. For GMP staining, sections were first stained with rat anti-mouse c-Kit (BioLegend, 135102) followed with a goat anti-rat-Cy3 (Jackson Immunoresearch, 112-165-167), and then with A488-conjugated lineage markers, Sca-1-A488 (BioLegend, 108116), CD150-A488 (BioLegend, 115916) and FcγR-A647 (BioLegend, 101314). After staining, all sections were counterstained with 1 μg/ml DAPI in PBS for 10 min at RT, mounted with Fluoromount G (Southern Biotech, 0100-01) and imaged on an SP5 upright confocal microscopes (Leica) with 20x objectives. For DGB staining of purified SECs, 2,000 cells were sorted directly onto poly-L-lysine coated slides (Sigma-Aldrich, P0425-72EA) and allowed to settle for 10 min before being fixed with 4% PFA for 10 min at RT, washed and counterstained with 1 μg/ml DAPI in PBS for 10 min at RT, mounted with VectaShield (Vector Laboratories, H-1200) and imaged on an A1 Ti Eclipse inverted microscope (Nikon) with 20x objectives. Images were processed using Volocity software (Perkin Elmer v.6.2) and analyzed with ImageJ. The find objects function in Volocity was used to quantify vascular volume based on Laminin staining.

### Cytokine profiling

For each individual mouse, BM fluid was flushed out from the four hind leg bones (two femurs and two tibiae) using the same 200 μl of HBSS/2% FBS in a 1 ml syringe with 26G needle. BM cells were sedimented by centrifugation at 300 g for 5 min and collected supernatants were purified by an additional centrifugation at 15,300 g for 10 min. BM fluids were stored at −80°C until use. For 200-plex cytokine array, BM fluid samples were submitted to Quantibody Testing Service (Raybiotech) and diluted 4x prior to analyses. For bead array analysis, 50 μl of 2x diluted sample was analyzed using Mouse 20-Plex panel (Thermo Fischer Scientific) using either a Magpix (UCSF) or Luminex 200 (CUIMC) analyzer according to manufacturer’s protocol. For ELISA measurements, 4x-diluted (SCF, SDF1a, TPO; Raybiotech) and 90x-diluted (TGF-β; R&D Systems) samples were prepared according to the manufacturer’s instructions and analyzed on a Synergy 2 plate reader (Biotek).

### Plate-based scRNAseq

Samples were processed following modifications to the Smart-Seq2 protocol^41^ based in the mcSCRB-Seq protocol^42^. Single endosteal and central marrow stromal cells were directly sorted into individual wells of a 96-well PCR plate in 2.3 μl of lysis buffer containing 0.2% Triton X-100 (Sigma-Aldrich) and 1U of Superase-In RNase Inhibitor (Ambion). Of note, information regarding expression of surface markers was recorded for each cell when sorting. Cells were frozen immediately at −80C until further processing. After thawing on ice, 2 μl of an annealing mixture containing1 μM oligodT (IDT), 5 mM each dNTPs and a 1:6,000,000 dilution of ERCC RNA Spike-In Mix (Invitrogen) was added followed by incubation at 72 °C for 3 min. Then 5.7 μl of a Reverse Transcription mix containing 3.5 U/μl of Maxima H minus retrotranscriptase (ThermoFisher), 0.88 U/μl of Superase-In RNase Inhibitor, 1.75x Maxima RT Buffer, 3.5 μM TSO (Qiagen) and 13.15% PEG 8000 (Sigma-Aldrich) was added and the mixture was incubated at 42 °C for 90 min, followed by 70 °C for 15 min. cDNA was further amplified by adding 40 μl of a PCR mix containing 0.03 U/μl of Terra PCR direct polymerase (Takara Bio), 1.25x Terra PCR Direct Buffer, and 0.25 μM IS PCR primer (ID). PCR was as follows: 3 min at 98 °C for initial denaturation followed by 21 cycles of 15 s at 98 °C, 30 s at 65 °C, 4 min at 68 °C. Final elongation was performed for 10 min at 72 °C. Sequences of oligodT, TSO and IS PCR primers were as previously described^41^. Following preamplification, all samples were purified using Ampure XP beads (Beckman Coulter) at a ratio of 1:0.6 with a final elution in 25 μl of EB Buffer (Qiagen). The cDNA was then quantified using the Quant-iT PicoGreen dsDNA Assay Kit (Thermo Fisher). Size distributions were checked on high-sensitivity DNA chips (Agilent Bioanalyzer). Samples were used to construct Nextera XT libraries (Illumina) from 100 pg of preamplified cDNA. Libraries were purified and size selected (0.5x-0.7x) using Ampure XP beads. Then, libraries were quantified using KAPA qPCR quantification kit (KAPA Biosystems), pooled and sequenced in an Illumina HiSeq 4000 instrument. Reads were mapped to the *Mus musculus* genome (EMSEMBL GRCm38.p4 Release 81) and ERCC sequences using GSNAP (version 2015-09-29) with parameters: −A sam –B 5 –t 24 –n 1 –Q –N 1. HTseq-count^43^ was used to count reads mapped to each gene, with parameters: –s no. All cells with < 100,000 reads mapping to endogenous RNA and >20% reads mapping to mitochondrial genes were considered low quality and removed from downstream analyses. Overall, 661cells (67.38%) passed our quality controls distributed as follows: Young marrow EC (SEC), 70; Old marrow EC (SEC), 71; Young endosteal EC (AEC), 84; Old endosteal EC (AEC), 82; Young MSC-L, 53; Old MSC-L, 58; Young MSC-S, 59; Old MSC-S, 59; Young mOPr, 26; Old mOPr, 26; Young OPr, 44; Old OPr, 29. The mean of nuclear reads mapped per cell was 471,244. Data were normalised and highly variable genes were identified as previously described^44^, using a false discovery rate threshold equal to 0.1 for the chi-squared test. Only highly variable genes were considered to perform PCA analysis, using the *prcomp* function in R (version 3.6.3). UMAP was calculated using *umap* function (version 0.2.7) in R based on normalised expression value with 15 nearest neighbors. All genes were used for ICGS clustering (AltAnalyzer)^45^ of mesenchymal and endothelial populations. In both cases we used version (v2.0) and ‘stringent’ with cell cycle and all other parameters as default. For mesenchymal populations, 16 clusters of cells (denoted M1 to M16) and 6 clusters of genes (denoted A to F) were defined. For endothelial populations, 7 clusters of cells (denoted E1 to E7) and 6 clusters of genes (denoted a to g) were defined. For subsequent analysis, mesenchymal clusters of cells were pooled as follows: M1 and M2 for “MSC-L-like”; M3, M4 and M5 for “OPr-like”; M6 to M16 for “MSC-S-like”. Subsequent endothelial groups of cells were defined as follows: clusters E3 and E4 constituted “AEC-like”; cluster E7 “SEC-like”. For MSC-L gene identity comparison, the geometric mean of the expression of all genes contained in cluster A was calculated for each cell of clusters M1 and M2 and then clusters M1 and M2 were compared using the independent two-sided t test (p-value < 2.2e-16). Differential expression analysis of old and young cells within each group was performed using DESeq2^46^ version 1.26.0. For violin plots representation, normalised expression values of *Kitl*, *Cxcl12*, *Il1a, Il1b* and *Il1rn* were plotted in various groups for young and old cells using the *ggplot2* package (version 3.3.0) in R with scale parameter set to ‘width’. Pathway analysis was conducted using Gene Set Enrichment Analysis (GSEA) software v4.1.0. Gene symbols were mapped to MSigDB.v7.2.chip and overlaps with Hallmark (h.all.v7.2.symbols.gmt) gene sets were determined using the classic scoring scheme. For endothelial cell types, no FDR<0.05 significant enrichments were obtained using GSEA, and so Ingenuity Pathway Analyses software (Qiagen, December 2020) was eventually employed. For IPA of endothelial cells, log 2 fold change of expression and p-adjusted values for all genes with nominal p values under 0.05 were imported into the software and IPA Canonical Pathways were determined.

### Droplet-based scRNAseq

For both endosteal and central marrow populations, 30,000-50,000 Ter119^−^/CD45^−^ stromal cells were sorted into 1.5 ml tubes containing 500 μl of filter-sterilized αMEM with 10% FBS and transferred to the Columbia Genome Center Single Cell Analysis Core for microfluidic cell processing, library preparation and sequencing. In brief, cells were re-counted and viability was assessed using a Countess II FL Automated Cell Counter (Thermo), and samples were processed following manufacturer’s recommendations for Chromium Single Cell 3′ Library & Gel Bead Kit v2 (10X Genomics). 17,500 cells were loaded for each sample and 1 sample was loaded per condition. Samples were sequenced in Illumina HiSeq4000 sequencer machine. We obtained an average of ~286 million reads per sample. The alignment was done using *Cellranger* (version 2.1.1). Top 2,300 barcodes were selected for samples corresponding to young and old central marrow and young endosteum. Top 2,858 barcodes were selected for the sample corresponding to old endosteum. The downstream analysis was done using *Scanpy*^47^ (version 1.6.0) in *Python* (version 3.7.1). For these samples, 757 doublets were estimated and removed using *Scrublet* package^48^ (version 0.2.1). Further quality control (QC) was performed based on 3 parameters: 1) cells with at least 200 and no more than 7,000 genes detected; 2) cells with less than 65,000 associated counts; 3) cells with less than 5.5% of UMI counts associated to mitochondrial genes. After QC, 8,735 cells were kept for subsequent analysis. In addition, only genes that have more than 1 UMI count in at least 5 cells were maintained in further analysis. Cells were then normalised to 10,000 UMIs per cell and logarithmically transformed. Highly variable genes (HVGs) were selected using “highly_variable_genes” method with “min_mean=0.0175, max_mean=3, min_disp=0.5”. Read depth, number of genes and number of mitochondrial counts were removed using the “regress_out” function in *Scanpy*. UMAP visualisations were obtained from 50 PCA components and 10 neighbors using *Scanpy*. Clusters were defined using *Louvain* clustering (version 0.7.0). Clusters containing stromal cells were subset based on the expression of genes expressed mainly in either hematopoietic cells or stromal cells. Only clusters that contained cells that expressed stromal marker genes were kept for subsequent analysis. In total, 2378 cells were selected, distributed as follows: 188 cells came from the young central marrow sample, 726 cells from the old central marrow sample; 1320 cells came from the young endosteum sample and 144 cells from the old endosteum sample. Highly variable genes (HVGs) were then obtained for the selected cells and the effects of read depth, number of genes and number of mitochondrial counts were regressed, as indicated above. The UMAP visualisation was obtained as before and again clusters were defined using Louvain clustering with “*resolution 0.3*”. Cell types were annotated using typical marker genes for the different populations.

### Quantitative RT-PCR analyses

10,000-50,000 cells per population were sorted directly into RPE buffer (Qiagen) and stored at −80°C until purification with the RNeasy Plus Mini Kit (Qiagen) according to manufacturer’s protocol. Following column purification, RNA was immediately reverse-transcribed using SuperScriptIII kit with random hexamers (Invitrogen). qPCR runs were performed on a QuantStudio 7 Flex Real-Time PCR System (Applied Biosystems) using SYBR Green reagents (Applied Biosystems) and the cDNA equivalent of 200 cells per reaction template. Sequences for qRT–PCR primers were: *Il1a*, forward GCACCTTACACCTACCAGAGT and reverse TGCAGGTCATTTAACCAAGTGG (NM_010554); *Il1b*, forward GCAACTGTTCCTGAACTCAACT and reverse ATCTTTTGGGGTCCGTCAACT (NM_008361); *Tnf*, forward AGGGATGAGAAGTTCCCAAAT and reverse GCTTGTCACTCGAATTTTGAG (NM_013693); and *Gapdh*, forward GACTTCAACAGCAACTCCCAC and reverse TCCACCACCCTGTTGCTGA (NM_008084). Cycle threshold values were normalized to *Gapdh*.

### Fluidigm analyses

Gene expression analyses using the Fluidigm 96.96 Dynamic Array IFC were performed as previously described^5^. Briefly, HSC, MPP3 and MPP4 (100 cells/well) were directly sorted per well of a 96-well plates containing 5 μl CellsDirect lysis buffer (Invitrogen, 11753-100), reverse-transcribed and pre-amplified for 18 cycles using SuperScript III Platinum Taq Mix (Invitrogen, 12574-026) with a custom-made set of 96 proprietary target-specific primers (Fluidigm). The resulting cDNA was analyzed on a Biomark system (Fluidigm) using EvaGreen SYBR dye (Bio-Rad, 172-5211). Data were collected with Biomark Data Collection Software (Fluidigm) and analyzed using Biomark qPCR software with a quality threshold of 0.65 and linear baseline correction. Melt curves and melting temperature values for each assay reaction were checked individually, and reactions with melt curves showing multiple peaks or poor quality were discarded. Results for 2 and 3 weeks post-transplantation data were recalculated with *Hprt* as housekeeping gene and genes not analyzed in one of three conditions were removed, leaving 64-68 genes excluding housekeeping genes (*Actb*, *Gapdh*, *Gusb* and *Hprt*) for further analyses and calculation of Pearson’s correlation coefficients. For each dataset, values for each gene were averaged, and fold change values were calculated against young non-transplanted control and converted to Log2 value. Similarity matrixes were visualized with *Morpheus* (Broad Institute).

### Microarray analyses

Microarray analyses of old HSCs, MPP3 and MPP4 were performed as previously described^5^. Three to five independent biological replicates were used for each population. Total RNA was isolated from 20,000 cells per population sorted directly into TRIzol-LS (Invitrogen) and purified using Arcturus PicoPure (Applied Biosystems) with RNase-free DNase (Qiagen). RNA was amplified, labeled, and fragmented using NuGEN Ovation Pico linear amplification kits (Nugen Technologies) and hybridized onto mouse Gene ST 1.0 arrays (Affymetrix). Gene expression microarray data were normalized using RMA followed by quantile normalization as implemented in the 2.15.1 R package (www.r-project.org) using a standard (lambda=1) exponential reference distribution. Significance Analysis of Microarrays (SAM)^49^ was then performed on young and old cells within a population to determine SAM delta scores. Pathway analysis was performed using GSEA (v4.1.0) on the top 1000 up and downregulated genes in young versus old for each population. Gene symbols were mapped to MSigDB.v7.2.chip and overlaps with Reactome (c2.cp.reactome.v7.2.symbols.gmt) gene sets were determined using the classic scoring scheme.

### Statistics

All experiments were repeated as indicated; n indicates the numbers of independent biological repeats. Data are expressed as mean ± standard deviation (S.D.) or standard error of the mean (S.E.M.) as indicated. Mice for treatment and transplantation were randomized, samples were alternated whenever possible, and no blinding protocol was used. No statistical method was used to predetermine sample size. Pairwise statistical significance was evaluated by two-tailed Student’s t-test. *P* values < 0.05 were considered statistically significant. Figures were made with GraphPad Prism software.

## Data availability

Data sets that support the findings of this study have been deposited in the Gene Expression Omnibus (pending accession numbers). Source data for all the figures are provided with the paper. All other data are available from the corresponding author upon reasonable request.

## Notes

### Competing Interest Statement

The authors have declared no competing interest.

