## Extended Data Figure 1-10 for "Stromal inflammation is a targetable driver of hematopoietic aging"

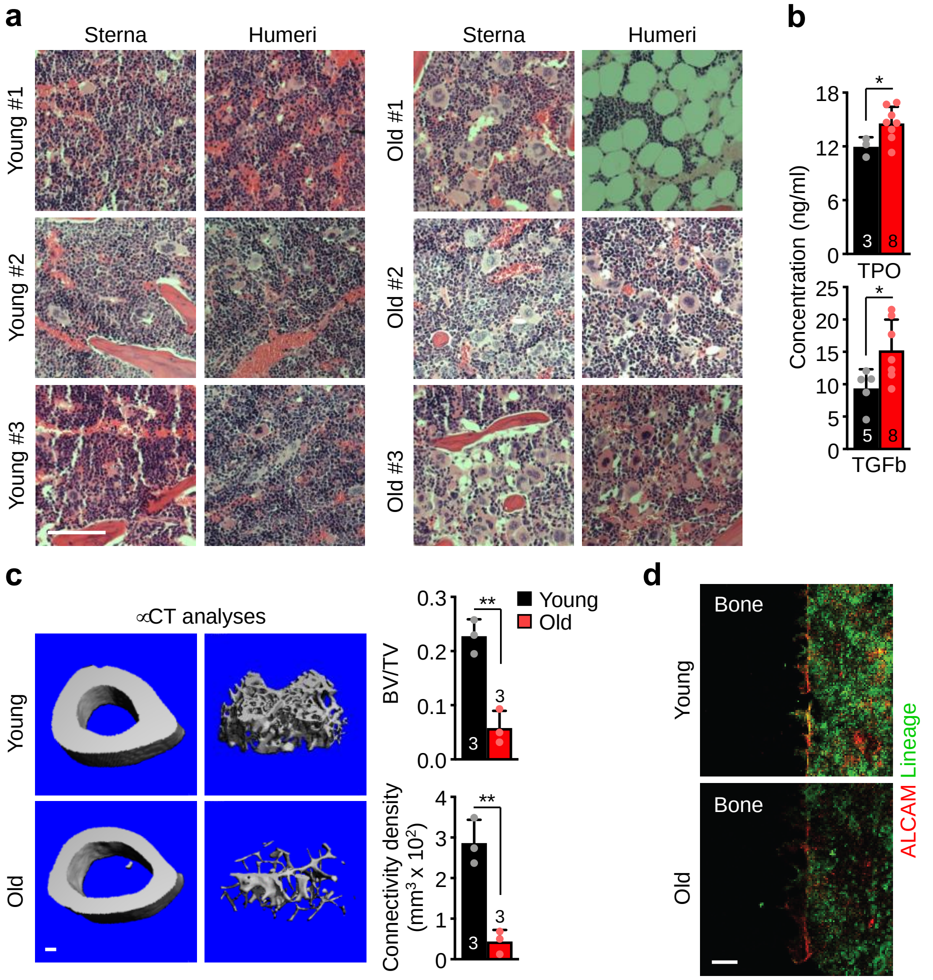


**Extended Data Figure 1 | Gross analysis of the remodeled old BM cavity. a**, H&E staining of humeri and sterna of 3 individual young and old mice. Scale bar, 100 µm. **b**, TPO and TGFβ levels in young and old BM fluids. **c**, µCT analyses of young and old femurs with representative images of cortical and trabecular regions (left) and quantification of bone volume/total volume (BV/TV) and connectivity density (right). **d**, Representative image of bone-lining ALCAM$^{+}$ osteoblasts immunofluorescence staining in young and old mice. Scale bar, 100 µm. Data are means ± S.D; *p ≤ 0.05, **p≤0.01.

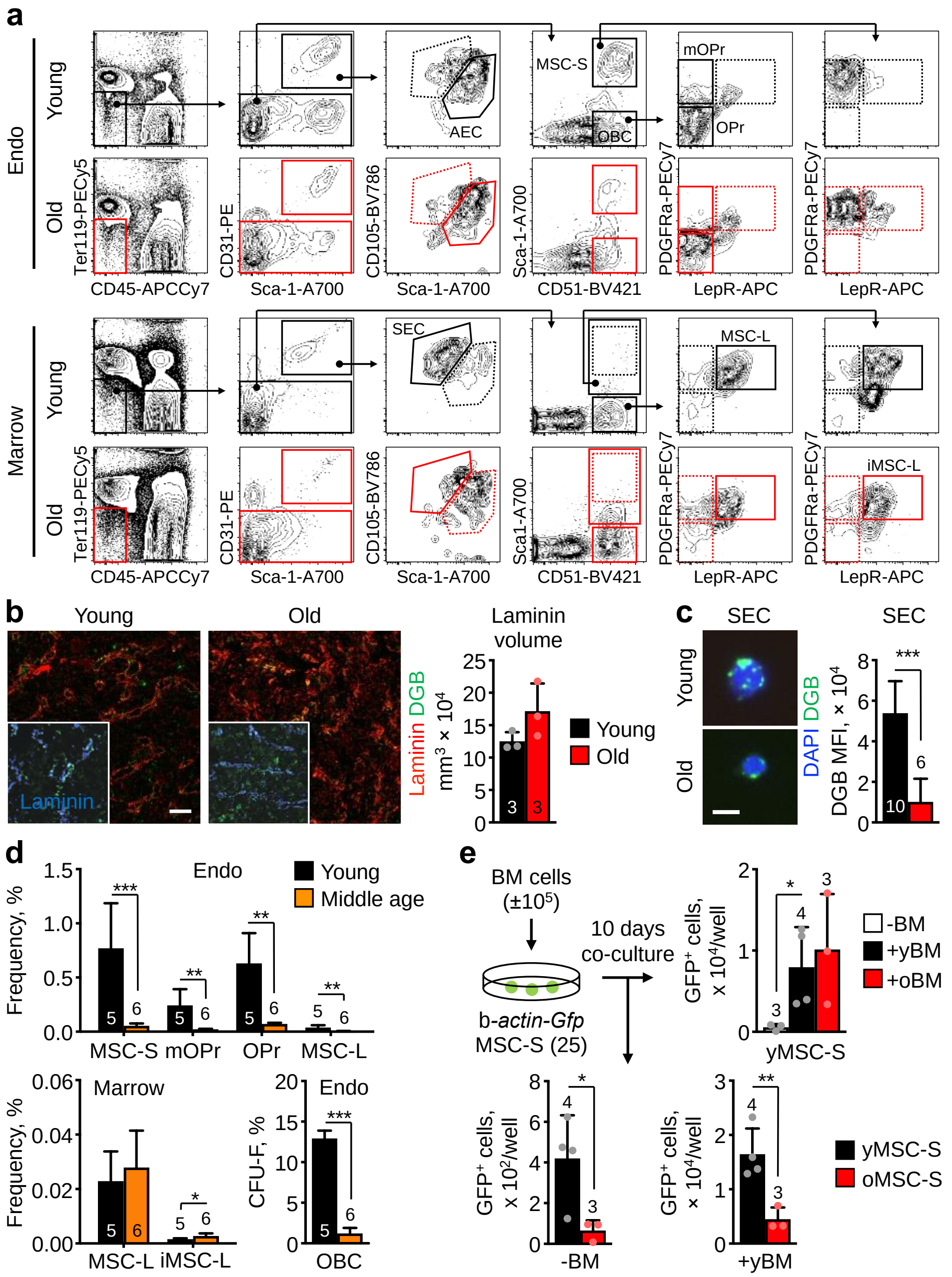


**Extended Data Figure 2 | Characterization of young and old BM niche cells. a**, Representative flow cytometry staining of young and old stromal populations. AEC, arteriolar endothelial cell; SEC, sinusoidal endothelial cell; MSC-S, Sca-1$^{+}$ mesenchymal stromal cell; mOPr, multipotent osteoprogenitor; OPr, osteoprogenitor; MSC-L, LepR$^{+}$ mesenchymal stromal cell; iMSC-L, inflammatory Sca-1$^{low}$ MSC-L. **b,** Representative images and quantification of immunofluorescence staining of vascular volume (laminin) and vascular leakage by dragon-green beads (DGB) diffusion assay in young and old BM. Scale bar, 50 µm. **c,** Representative images and quantification by flow cytometry of DGB endocytosis in young and old marrow SEC. Scale bar, 5 µm. **d**, Frequency of endosteal and marrow mesenchymal populations in young and middle age (13 month-old) mice with changes in CFU-F from endosteal OBCs (bottom right)**. e**, Experimental scheme for the indicated co-culture experiments showing the effects of young or old BM cells on young MSC-S (top right), and young BM cells on young and old MSC-S (bottom). Data are means ± S.D; *p ≤ 0.05, **p ≤ 0.01, ***p ≤ 0.001.

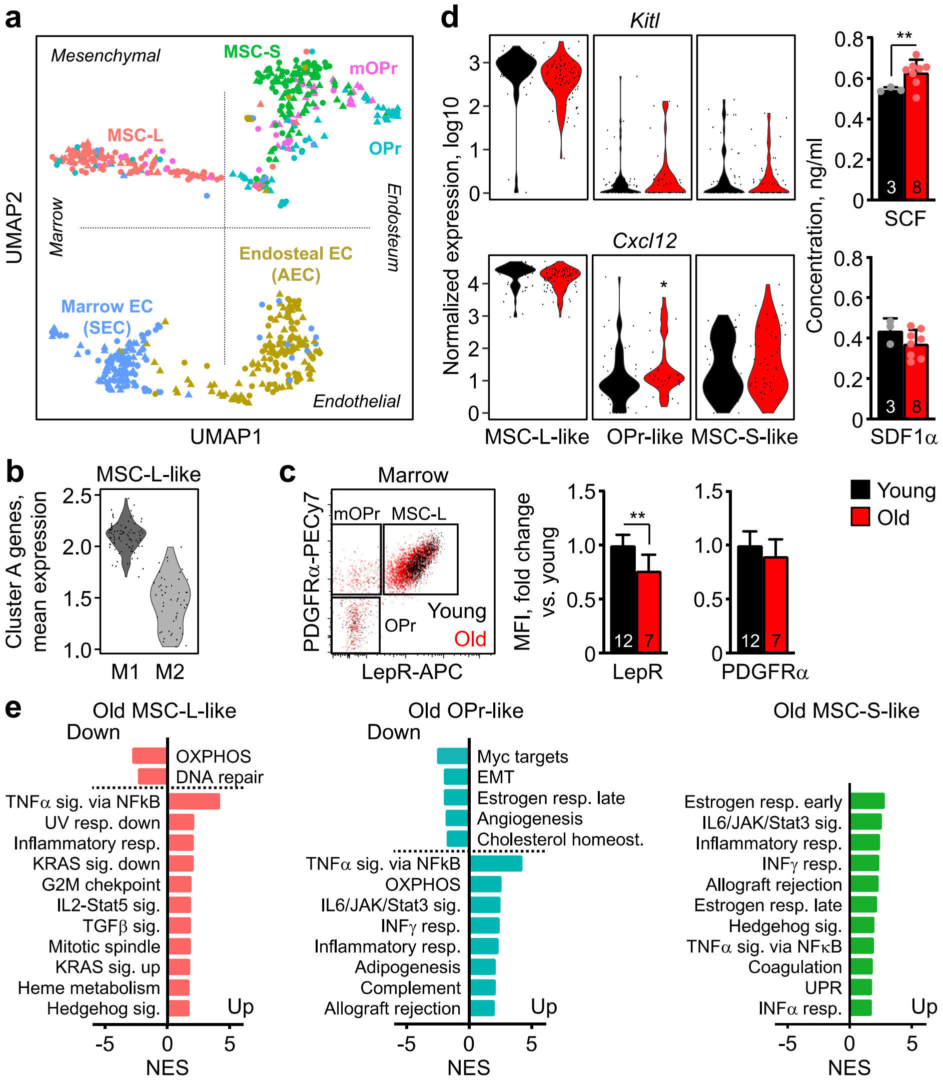


**Extended Data Figure 3 | Molecular features of old mesenchymal populations. a,** UMAP visualization of the entire plate-based scRNAseq dataset. **b**, Global changes in MSC-L gene identity in clusters M2 vs. M1. **c**, Representative flow cytometry staining (top) and quantification of LepR and PDGF-Rα levels (bottom) in young and old MSC-L. **d**, HSC niche factors with violin plots representation of *Kitl* and *Cxcl12* expression in the indicated young and old mesenchymal populations (left; *padj ≤ 0.05) and SCF and SDF1α levels in young and old BM fluids (right). **e,** GSEA results for Hallmark biological processes significantly affected in old MSC-L-like, OPr-like and MSC-S-like groups. Data are means ± S.D except for results from the plate-based scRNAseq dataset shown in b and d; *p ≤ 0.05, **p ≤ 0.01.

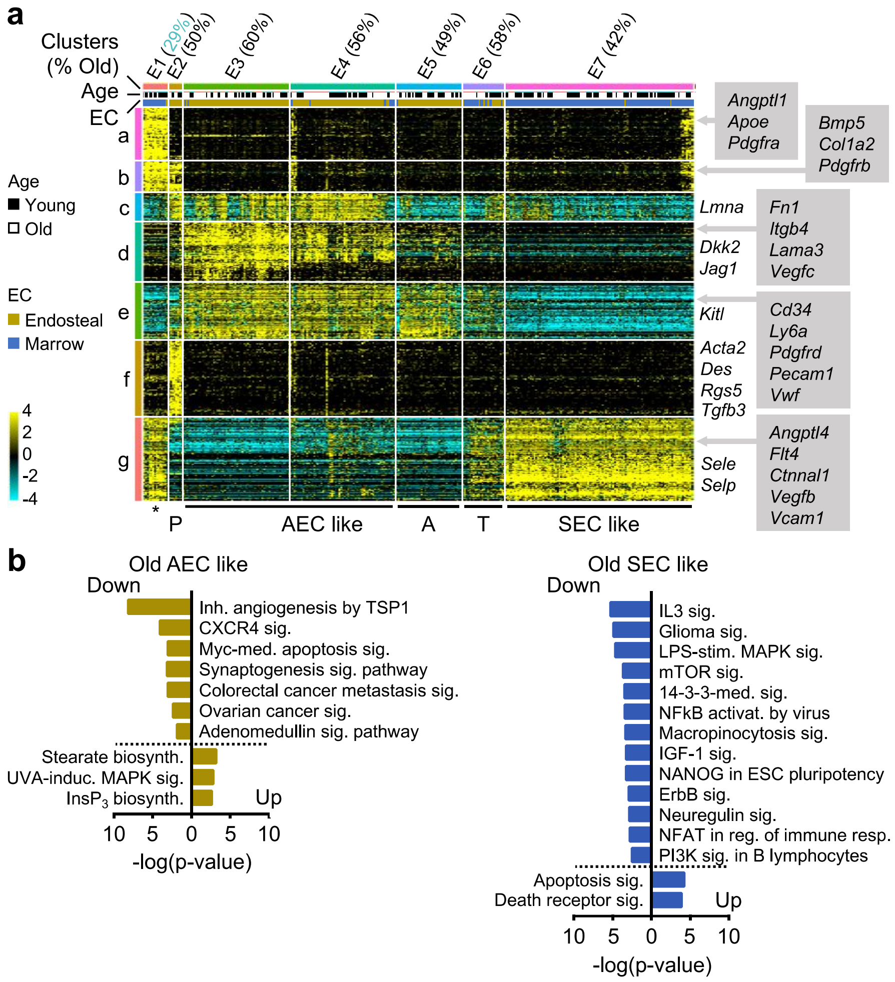


**Extended Data Figure 4 | Molecular features of old endothelial populations. a,** ICGS output of young and old endothelial populations with 7 clusters of cells (E1 to E7) defined according to the expressing pattern of the 7 clusters of genes (a to g). Examples of genes included in gene clusters a to g are shown. Star, contaminating mesenchymal/endothelial doublets; P, pericytes; A, arteriols; T, transition vessels. **b**, Ingenuity Pathway Analysis (IPA) canonical pathways enriched in old AEC-like and SEC-like groups. All results are from the plate-based scRNAseq dataset.

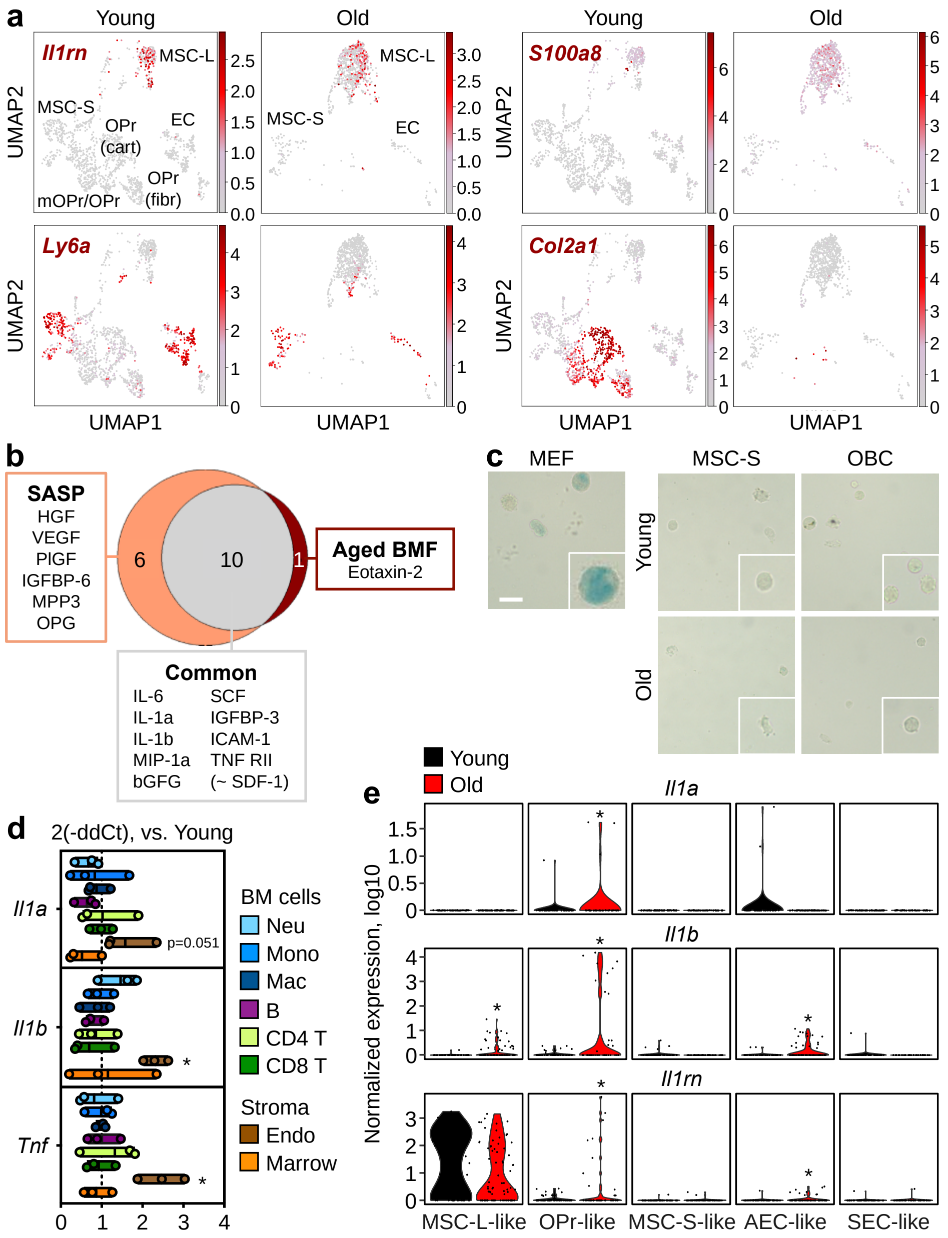


**Extended Data Figure 5 | Details of the inflammatory old BM niche. a,** characteristic expression patterns for the indicated genes in the droplet-based scRNAseq dataset. Cells in the UMAP were colored according to the expression levels of the indicated genes. Color scheme is based on ln scale of normalized counts from 0 (gray) to the indicated maximum value in the scale (dark red). **b**, Overlap between cytokines upregulated in old BM fluids and published SASP profile. **c**, Representative SA-β-gal staining in control irradiated mouse embryonic fibroblasts (MEF) and isolated young and old MSC-S and OBC. Scale bar, 20 µm. **d**, qRT-PCR-based analyses of *Il1a*, *Il1b* and *Tnf* expression in the indicated BM and unfractionated endosteal and central marrow Ter119$^-$/CD45$^-$ stromaf fractions. Neu, CD3$^-$/B220$^-$/NK1.1$^-$/Mac-1$^+$/Ly-6G$^+$/Ly-6C$^{mid}$ neutrophil; Mono, CD3$^-$/B220$^-$/NK1.1$^-$/Mac-1$^+$/Ly-6G$^-$/Ly-6C$^{hi}$ monocyte; Mac, CD3$^-$/B220$^-$/NK1.1$^-$/Mac-1$^+$/Ly-6G$^-$/Ly-6C$^-$ macrophage; B, B220$^+$/CD19$^+$ B cell; CD4 T, CD4$^+$/TCRβ$^+$ T cell; CD8 T, CD8$^+$/TCRβ$^+$ T cell. Results are expressed as box plot with mean and minimal/maximal values; *$p \leq 0.05$. **d**, Violin plots representation of *Il1a*, *Il1b*, and *Il1rn* expression in the indicated young and old mesenchymal and endothelial like groups. Results are from the plate-based scRNAseq dataset; *padj ≤ 0.05.

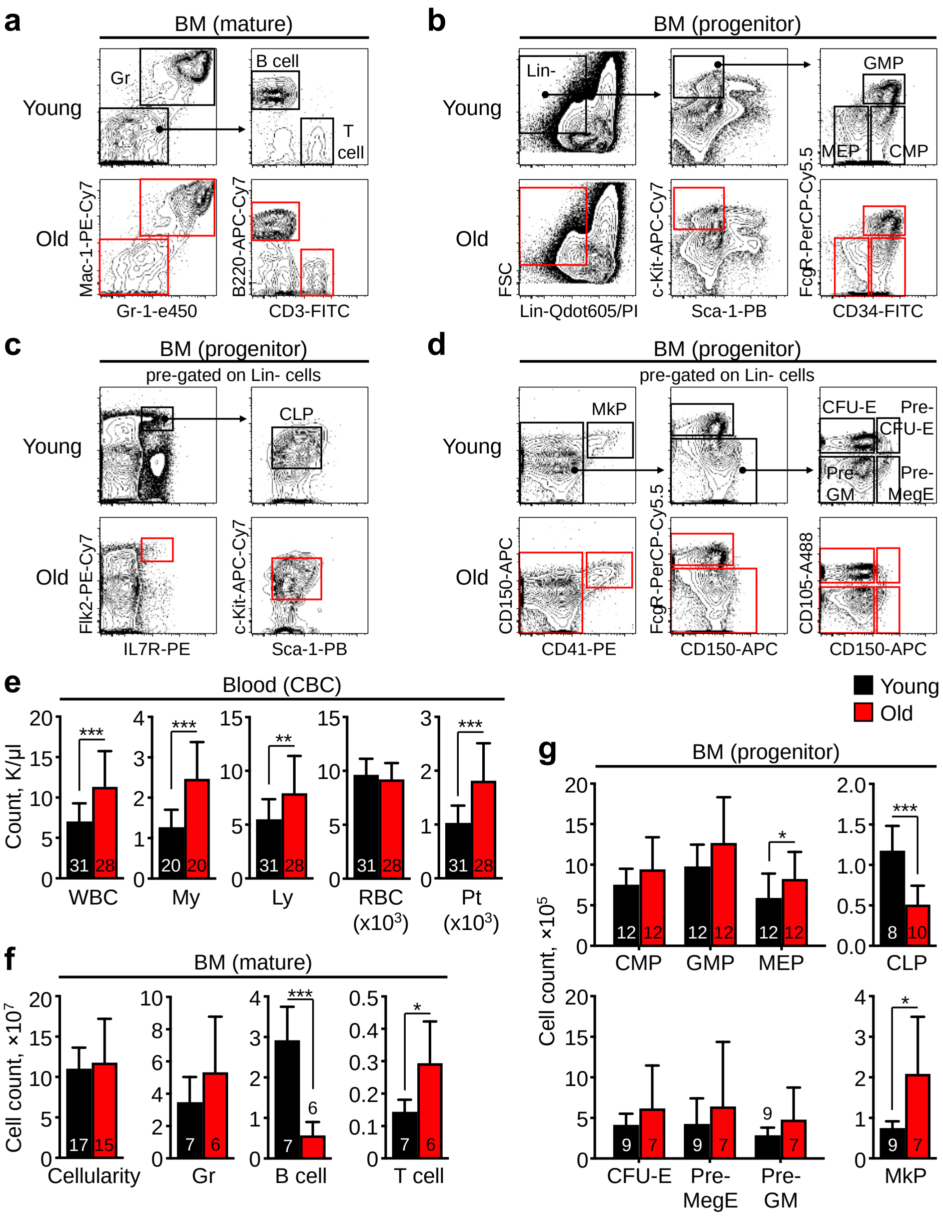

**Extended Data Figure 6 | Age-related changes in the blood and BM. a-b**, Representative flow cytometry staining of young and old BM cells with gating strategy for (a) granulocytes (Gr), B and T cells, (b) myeloid progenitors, (c) lymphoid progenitors, and (d) Mk and erythroid progenitors. CMP, common myeloid progenitor; GMP, granulocyte-macrophage progenitor; MEP, megakaryocyte-erythrocyte progenitor; CLP, common lymphoid progenitor; MkP, megakaryocyte progenitor; Pre-GM, pre-granulocyte/macrophage; Pre-MegE, pre-megakaryocyte/erythrocyte; CFU-E, erythroid colony-forming unit. **e,** Complete blood count (CBC) parameters in young and old mice. WBC, white blood cell; My, myeloid (neutrophil + basophil + eosinophil); Ly, lymphocyte; RBC, red blood cell; Pt, platelet. **f,** Cellularity and quantification of mature populations in young and old BM. **g**, Quantification of progenitor populations in young and old BM. Data are means ± S.D; *p ≤ 0.05, **p ≤ 0.01, ***p ≤ 0.001.

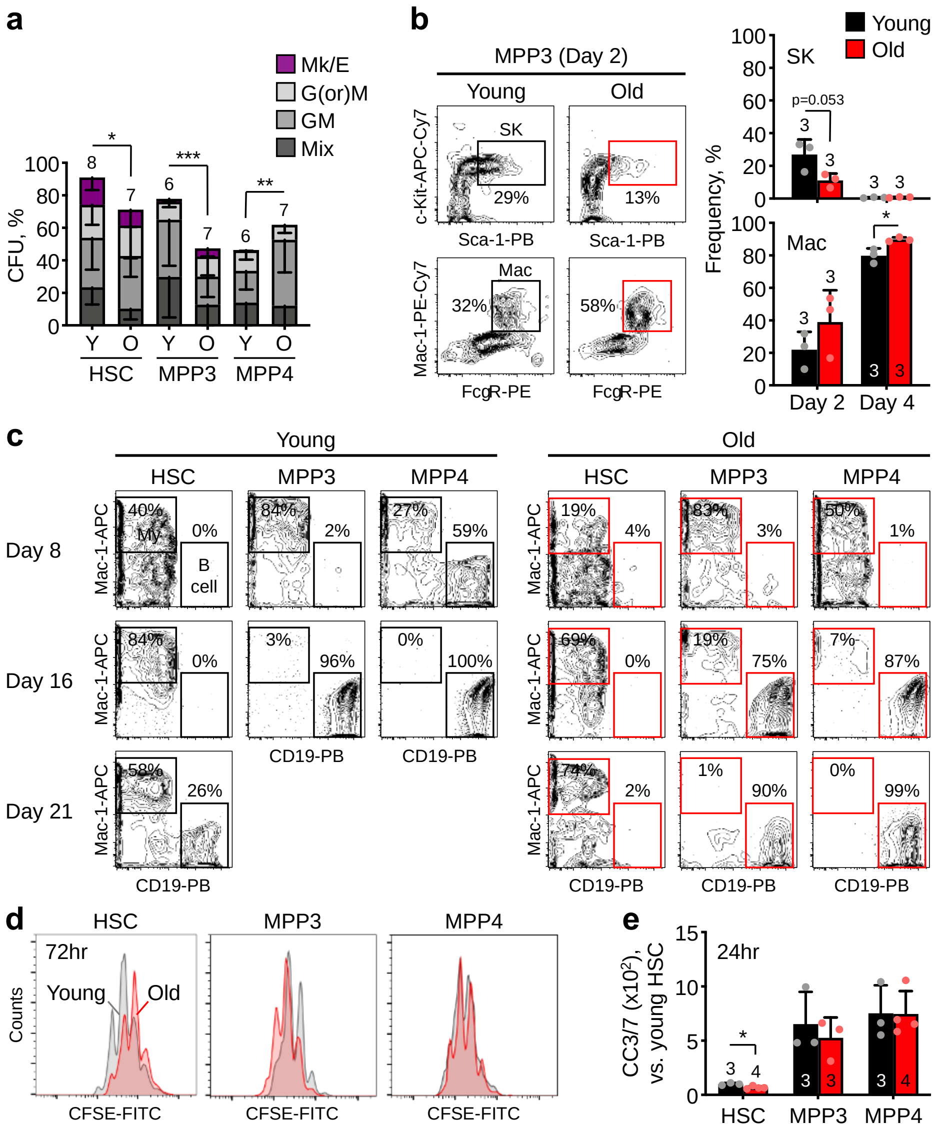


**Extended Data Figure 7 | Altered functionality of old HSCs and MPP populations. a**, Colony formation in methylcellulose for young and old HSCs, MPP3 and MPP4. Mix: all lineages; GM: granulocyte/macrophage; G(or)M: granulocyte (or) macrophage; MegE: megakaryocyte/erythrocyte; CFU: colony-forming units. **b**, Myeloid differentiation in liquid culture for young and old MPP3 with representative flow cytometry staining (left) and quantification (right) of immature Sca-1$^{+}$/c-Kit$^{+}$ (SK) and mature Mac-1$^{+}$/FcγR$^{+}$ macrophage (Mac). **c,** Representative flow cytometry staining of CD19$^{+}$ lymphoid vs. Mac-1$^{+}$ myeloid differentiation in OP9+IL7 culture conditions for young and old HSCs, MPP3 and MPP4. Results are representative of 3 independent experiments. **d**, Representative histograms of CFSE staining of cultured young and old HSCs, MPP3 and MPP4. Results are representative of 3 independent experiments. **e**, Cleaved caspase 3/7 (CC3/7) activity in cultured young and old HSCs, MPP3 and MPP4. Data are means ± S.D; *p ≤ 0.05, **p ≤ 0.01, ***p ≤ 0.001.

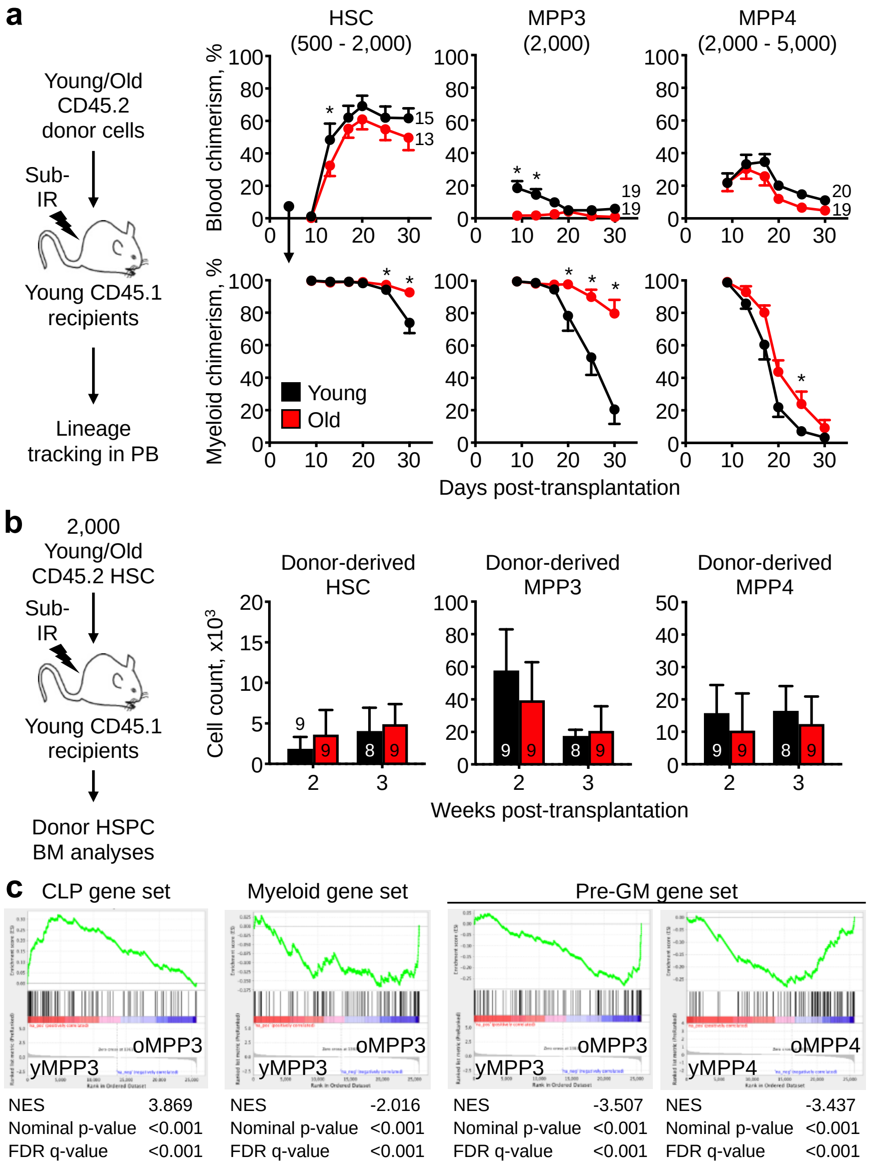


**Extended Data Figure 8 | Lineage biases in old MPPs. a**, Short-term lineage tracking following transplantations of young and old HSCs, MPP3 and MPP4 in sub-lethally irradiated recipients with experimental scheme (left) and quantification of overall blood donor chimerism (top graphs) and myeloid chimerism among donor cells (bottom graphs). Data are means ± S.E.M. **b**, Short-term regeneration of donor HSC, MPP3 and MPP4 compartment following transplantations of young and old HSCs in sub-lethally irradiated recipients with experimental scheme (left) and quantification of donor-derived populations (right). **c,** GSEA for lineage differentiation genes in microarray analyses of young and old MPP3 and MPP4 (Myeloid genes, GSE6506; CLP and pre-GM genes, GSE8407). Data are means ± S.D. except when indicated; $*p \leq 0.05$, $**p \leq 0.01$, $***p \leq 0.001$.

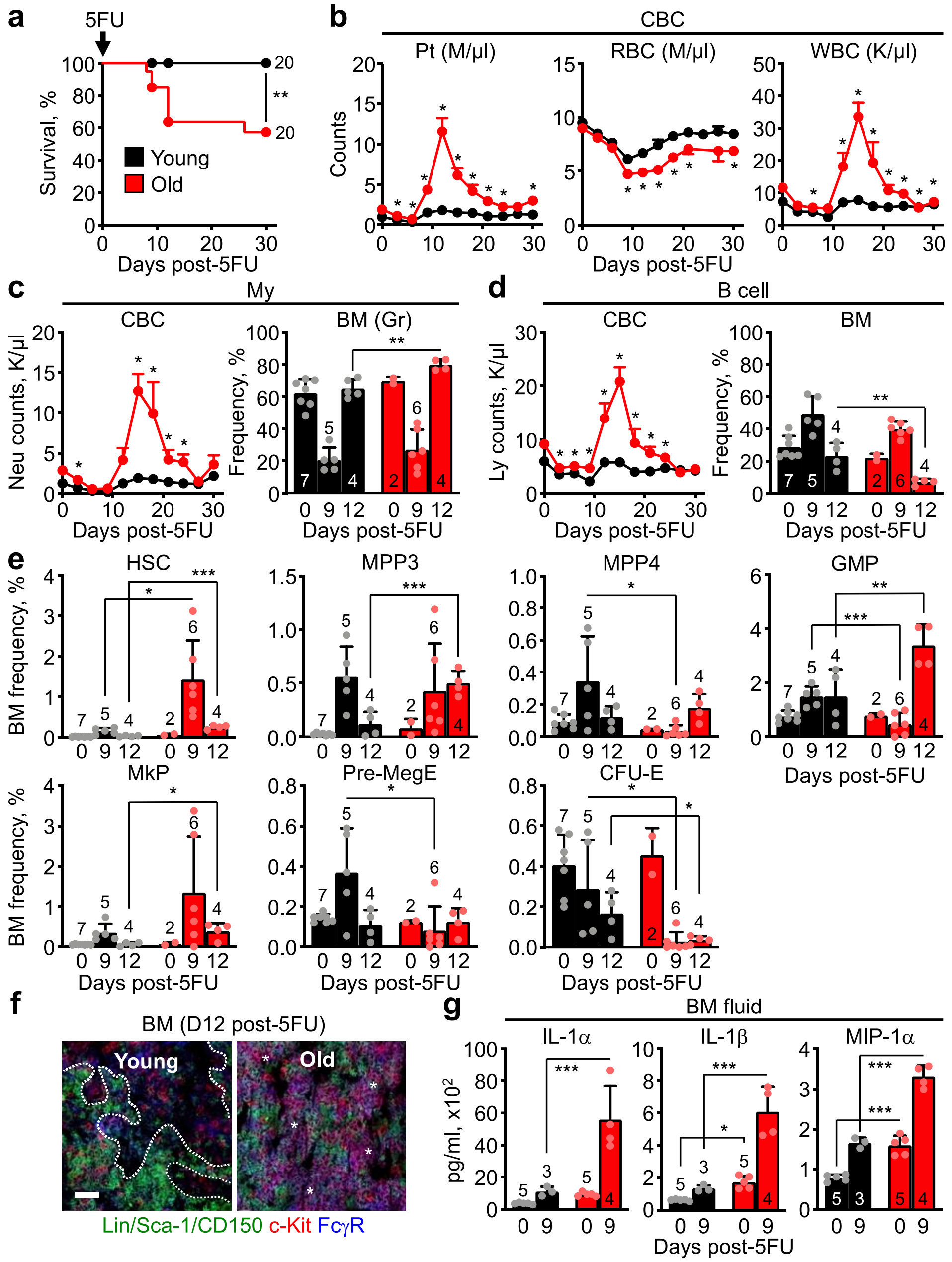


**Extended Data Figure 9 | Impaired hematopoietic regeneration in old mice**. **a,** Experimental scheme and survival of young and old mice following 5FU injection. **b**, Blood regeneration post-5FU treatment in young and old mice. For all CBC data, each group was started with 20 individual mice and results are mean ± S.E.M. **c-d**, Regeneration of (c) myeloid and (d) B cell populations post-5FU treatment of young and old mice with quantification of changes in the blood (left) and BM (right). **e,** Regeneration of the indicated BM progenitor populations post-5FU treatment of young and old mice. **f**, Representative image of GMP immunofluorescence staining post-5FU treatment of young and old mice. Dotted lines indicate GPM clusters and stars GMP patches. Scale bar, 60 μm. **g**, Changes in IL-1α and IL-1β levels in BM fluids post-5FU treatment of young and old mice. Data are means ± S.D. except when indicated; *p ≤ 0.05, **p ≤ 0.01.

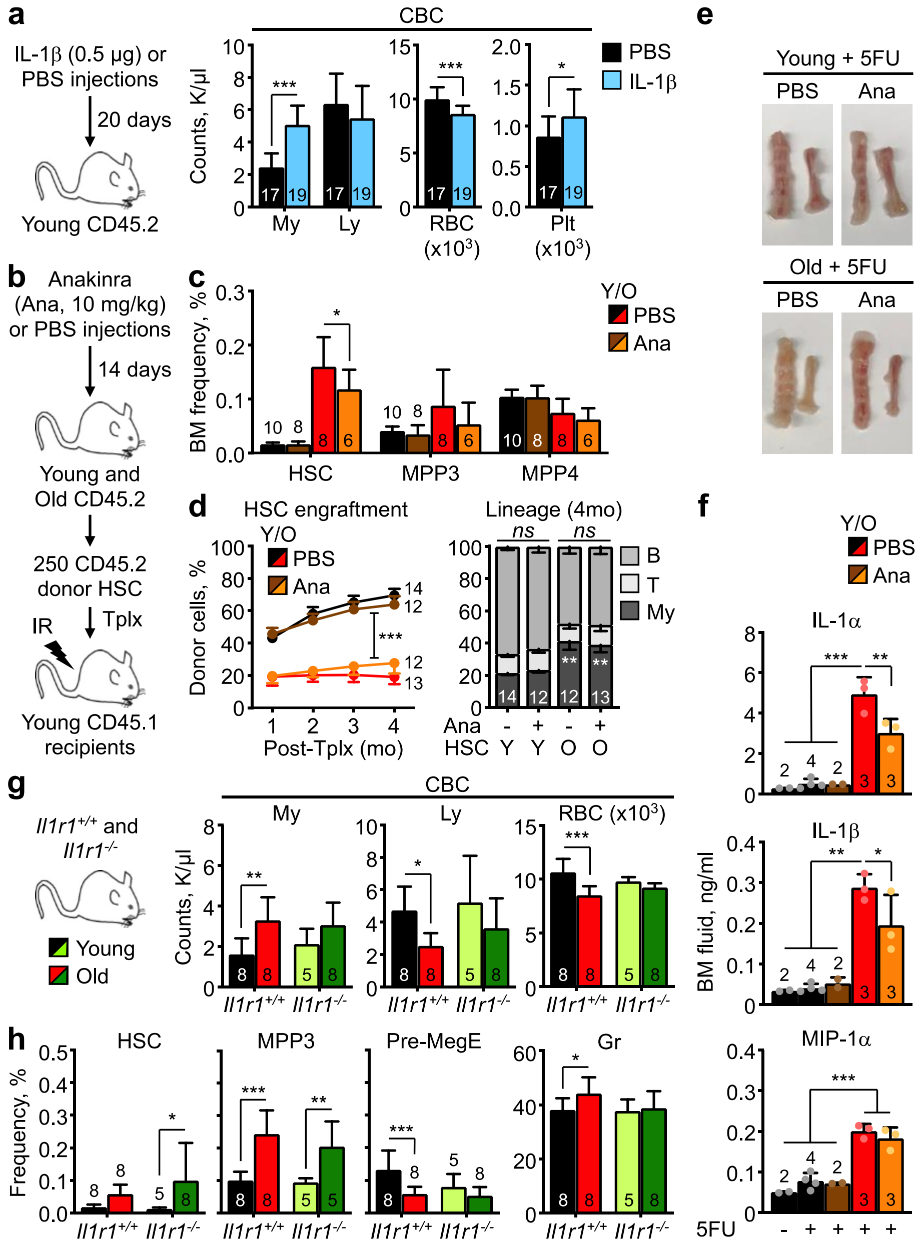


**Extended Data Figure 10 | Modulating IL-1 levels in young and old mice affect aging parameters.** **a,** Changes in blood parameters following chronic IL-1β injections in young mice with experimental scheme (left) and CBC values (right). **b-d,** Short-term blockade of IL-1 signaling upon anakinra (Ana) treatment in young and old mice with: (b) experimental scheme; (c) changes in HSCs and MPP frequency; and (d) engraftment over time (left, results are mean ± S.E.M.) and lineage reconstitution (right) at 4 months (4 mo) post-transplantation (Tplx) of the indicated HSC populations. **e-f,** Additional characterization of the effects of anakinra blockade of IL-1 signaling during 5FU-mediated regeneration in young and old mice with: (e) representative pictures of sterna and humeri; and (f) changes in IL-1α, IL-1β and MIP1α levels in BM fluids. **g-h**, Additional characterization of the effects of life-long loss of IL-1 signaling in young and old *Il1r1*$^{-/-}$ mice compared to age-matched *Il1r*$^{+/+}$ controls with changes in (g) blood parameters and (h) the indicated BM population frequency. Data are means ± S.D. except when indicated; *p ≤ 0.05, **p ≤ 0.01, ***p ≤ 0.001.
