## Supplementary Information Table 1-4 for "Stromal inflammation is a targetable driver of hematopoietic aging"

| | | |
|---|---|---|
| Under detection limit in both young and old | | |
| 4-1BB | I-TAC | Marapsin |
| 6Ckine | IFNg | MCP-1 |
| Activin A | IFNg R1 | MCP-5 |
| ADAMTS1 | IGFBP-2 | MCSF |
| ALK-1 | IL-1 R4 | MIP-1a |
| ANG-3 | IL-10 | MIP-1b |
| ANGPTL3 | IL-12p40 | MIP-3a |
| AR | IL-12p70 | MMP-10 |
| Artemin | IL-13 | OX40 Ligand |
| B7-1 | IL-15 | P-Cadherin |
| BLC | IL-17 | Pentraxin 3 |
| BTC | IL-17B | Persephin |
| CCL28 | IL-17B R | Prolactin |
| CD27 | IL-17E | Prostasin |
| CD27L | IL-17F | RAGE |
| CD30 | IL-1b | SCF |
| CD30L | IL-2 | SDF-1a |
| Chordin | IL-20 | sFRP-3 |
| CT-1 | IL-21 | Shh-N |
| CTLA4 | IL-22 | SLAM |
| DAN | IL-23 | TARC |
| DLL4 | IL-3 | TECK |
| EDAR | IL-3 Rb | Testican 3 |
| EGF | IL-33 | TGFb1 |
| Endocan | IL-4 | TIM-1 |
| Epigen | IL-5 | TNFa |
| Epiregulin | IL-6 | TPO |
| Fas | IL-7 | TRANCE |
| Fas L | IL-7 Ra | Tryptase ε |
| G-CSF | IL-9 | TSLP |
| Galectin-7 | Kremen-1 | TWEAK |
| GITR L | Leptin | VEGF |
| GM-CSF | Limitin | VEGF R2 |
| Gremlin | Lungkine | VEGF R3 |
| HAI-1 | Lymphotactin | VEGF-B |
| HGF R | MadCAM-1 | VEGF-D |
| Detected in all young but not old samples | | |
| Eotaxin-2 | IL-2 Ra | H60 |
| TCA-3 | Meteorin | TRAIL |
| TWEAK R | PlGF-2 | |
| Detected in all old but not young samples | | |
| Dtk | CD6 | GITR |
| CD36 | KC | |
| Above highest standards | | |
| Clusterin | IGF-1 | Periostin |

| | | |
|---|---|---|
| Cystatin C | L-Selectin | TCK-1 |
| Galectin-1 | MBL-2 | TNF RI |
| Galectin-3 | P-selectin | VCAM-1 |
| Not significantly changed | | |
| BAFF R | HGF | Neprilysin |
| CD40 | IL-28 | NOV |
| CD48 | Leptin R | OPN |
| CRP | Lipocalin-2 | PF4 |
| CXCL16 | LOX-1 | Pro-MMP-9 |
| Decorin | MFG-E8 | Progranulin |
| Fetuin A | MIG | Renin 1 |
| Fractalkine | MIP-2 | TREM-1 |
| gp130 | MIP-3b | TROY |
| Granzyme B | MMP-3 | |
| Significantly changed | | |
| Analyte | Young | Old |
| Adiponectin | 7593.4 ± 523.3 | 10644.5 ± 904.8*** |
| bFGF | 1091.3 ± 231.5 | 2164.7 ± 327.5*** |
| CCL6 | 6374.5 ± 337.8 | 4067.8 ± 446.8*** |
| Fcg RIIB | 10529.8 ± 2496.2 | 19569.5 ± 2019.8*** |
| IL-1a | 48.1 ± 6.3 | 95.4 ± 4.8*** |
| JAM-A | 3644.4 ± 750.9 | 8718.4 ± 768.8*** |
| Chemerin | 58082.5 ± 5765.8 | 38676.6 ± 10073.9** |
| Dkk-1 | 3668 ± 1070 | 1283.6 ± 227.1** |
| Eotaxin | 29.5 ± 9.6 | 64 ± 18.4** |
| Flt-3L | 3257.5 ± 535.4 | 4710 ± 527** |
| IGFBP-5 | 8607 ± 685.6 | 6690 ± 968.3** |
| IGFBP-6 | 2106.5 ± 283.1 | 1351.6 ± 272** |
| LIX | 804.8 ± 198.8 | 1746.3 ± 324** |
| RANTES | 82.1 ± 23.7 | 585.6 ± 171.3** |
| Resistin | 1860.6 ± 672.4 | 4302.7 ± 1217.3** |
| TACI | 10599.4 ± 12260.6 | 157233.9 ± 71367.6** |
| TNF RII | 2699.9 ± 342.1 | 3440.6 ± 216.7** |
| TremL1 | 32227 ± 3780 | 45409.9 ± 4734.9** |
| VEGF R1 | 2532 ± 592.1 | 5968.2 ± 1345.4** |
| ACE | 32359.4 ± 6618.2 | 43553.3 ± 7423.8* |
| C5a | 78.8 ± 14.7 | 107 ± 20.9* |
| CD40L | 1026.9 ± 357.2 | 1535.7 ± 237.8* |
| E-Cadherin | 8003.6 ± 1330.2 | 10267.8 ± 980.8* |
| E-selectin | 2207.3 ± 522.1 | 3372.1 ± 651.5* |
| Endoglin | 3484.8 ± 1484.7 | 7774.9 ± 2752.3* |
| Gas 1 | 1657.7 ± 385.3 | 2367.8 ± 307.8* |
| Gas 6 | 1952.9 ± 495.7 | 3482.9 ± 846.8* |
| ICAM-1 | 5172 ± 1733.6 | 8071.4 ± 1319.4* |
| IGFBP-3 | 13633.4 ± 3189 | 20585.3 ± 4191.9* |
| IL-1ra | 741.2 ± 475.1 | 4132.3 ± 1804.1* |
| MDC | 20.3 ± 3.5 | 11.5 ± 4.9* |
| MIP-1g | 1722.3 ± 154 | 2016.7 ± 202.1* |
| MMP-2 | 3568.7 ± 829.9 | 5844.7 ± 1645.2* |
| Nope | 1596.3 ± 148.6 | 1183.9 ± 306.9* |

| | | |
|---|---|---|
| OPG | 2212.2 ± 525.4 | 1299.2 ± 331.9* |
| Osteoactivin | 545.4 ± 224.6 | 3849.7 ± 1711.6* |
| PDGF-AA | 86.4 ± 27.1 | 197.5 ± 69.8* |

**SI Table 1 | Quantibody array-based measurements of cytokine concentration in young and old BM fluids.** BM fluids isolated from young (10 wks) and aged (27-29 mo) mice (n=5) was analyzed using Quantibody Testing Service (Raybiotech). Samples were diluted 4-fold prior to analyses. Results are mean ± S.D. and are expressed as pg/ml concentration; $*p \leq 0.05$, $**p \leq 0.05$, $***p \leq 0.05$.

**SI Table 2 | ICGS of young and old niche cells.** Smart-Seq2 gene expression data of mesenchymal and endothelial cells were separated and subjected to unsupervised single-cell population identification using Iterative Clustering and Guide-gene Selection (ICGS). The relative expression of every guide gene in each cluster, as well as the relative expression of every guide gene in each pooled group of clusters corresponding to the indicated cell types, was calculated by averaging the relative expression of each gene across individual cells within a cluster or group.

**SI Table 3 | DEGs and pathway analyses of young versus old niche cells.** Smart-Seq2 gene expression data between young and old cells in pooled identity groups as defined by ICGS were analyzed for differentially expressed genes (DEG) using the DESeq2 package. Overlap with Hallmark gene sets was evaluated, and statistically significant (FDR<0.05) overlaps are indicated in bold for mesenchymal populations. Since no significant overlaps were observed for endothelial populations, Ingenuity Pathway Analysis software (Qiagen) was employed to determine IPA Canonical Pathways with absolute z-score ≥ 1 and $-\log_{10}$(p-value) ≥ 2 enriched in young and old AEC-like and SEC-like cells.

**SI Table 4 | DEGs and pathway analyses of young versus old HSCs, MPP3 and MPP4.** Significance Analysis of Microarrays (SAM) was performed on young and old cells within each population to determine SAM delta scores. The top 1000 most highly DEGs for each group were collected. SAM scores for these genes were used for GSEA and overlap with Reactome gene sets was evaluated for each population. The top 5 statistically significant (FDR<0.05) overlaps for each group are in bold.
